# A unified genealogy of modern and ancient genomes

**DOI:** 10.1101/2021.02.16.431497

**Authors:** Anthony Wilder Wohns, Yan Wong, Ben Jeffery, Ali Akbari, Swapan Mallick, Ron Pinhasi, Nick Patterson, David Reich, Jerome Kelleher, Gil McVean

## Abstract

The sequencing of modern and ancient genomes from around the world has revolutionised our understanding of human history and evolution^1,2^. However, the general problem of how best to characterise the full complexity of ancestral relationships from the totality of human genomic variation remains unsolved. Patterns of variation in each data set are typically analysed independently, and often using parametric models or data reduction techniques that cannot capture the full complexity of human ancestry^3,4^. Moreover, variation in sequencing technology^5,6^, data quality^7^ and in silico processing^8,9^, coupled with complexities of data scale^10^, limit the ability to integrate data sources. Here, we introduce a non-parametric approach to inferring human genealogical history that overcomes many of these challenges and enables us to build the largest genealogy of both modern and ancient humans yet constructed. The genealogy provides a lossless and compact representation of multiple datasets, addresses the challenges of missing and erroneous data, and benefits from using ancient samples to constrain and date relationships. Using simulations and empirical analyses, we demonstrate the power of the method to recover relationships between individuals and populations, as well as to identify descendants of ancient samples. Finally, we show how applying a simple non-parametric estimator of ancestor geographical location to the inferred genealogy recapitulates key events in human history. Our results demonstrate that whole-genome genealogies are a powerful means of synthesising genetic data and provide rich insights into human evolution.

## Main

Our ability to determine relationships among individuals, populations and species is being transformed by population-scale biobanks of medical samples^11,12^, collections of thousands of ancient genomes^2^, and efforts to sequence millions of eukaryotic species^13^. Such relationships, and the resulting distributions of genetic and phenotypic variation, reflect the complex set of selective, demographic and molecular processes and events that have shaped species and are consequently a rich source of information about them^1,9,14,15^.

However, our ability to learn about evolutionary events and processes from the totality of genomic variation, in humans or other species, is limited by multiple factors. First, combining information from multiple data sets, even within a species, is technically challenging; discrepancies between cohorts due to error^16^, differing sequencing techniques^5,6^ and variant processing^8^ lead to noise that can easily obscure genuine signal. Second, few tools can cope with the vast data sets that arise from the combination of multiple sources^10^. Third, statistical analysis typically relies on data reduction techniques^17,18^ or the fitting of parametric models^4,19–21^, which will inevitably provide an incomplete picture of the complexities of evolutionary history. Finally, data access and governance restrictions often limit the ability to combine data sources^22^.

*Tree sequences* represent a potential solution to many of these problems^10,23^. Phylogenetic trees are fundamental to the evolutionary analysis of species; tree sequences extend this concept to multiple correlated trees along the genome, necessary when considering genealogies within recombining organisms^24^. Importantly, the tree sequence, and the mapping of mutation events to it, reflects the totality of what is knowable about genealogical relationships and the evolutionary history of individual variants. Fig. 1a shows how the tree sequence is defined as a graph with a set of nodes representing sampled chromosomes and ancestral haplotypes, edges connecting nodes representing lines of descent, and variable sites containing one or more mutations mapped onto the edges. Recombination events in the ancestral history of the sample create different edges and thus distinct, but highly correlated trees along the genome. Tree sequences contain a comprehensive record of ancestral history and can not only be used to compress genetic data^10^, but also lead to highly efficient algorithms for calculating population genetic statistics^25^.

**Fig. 1:**
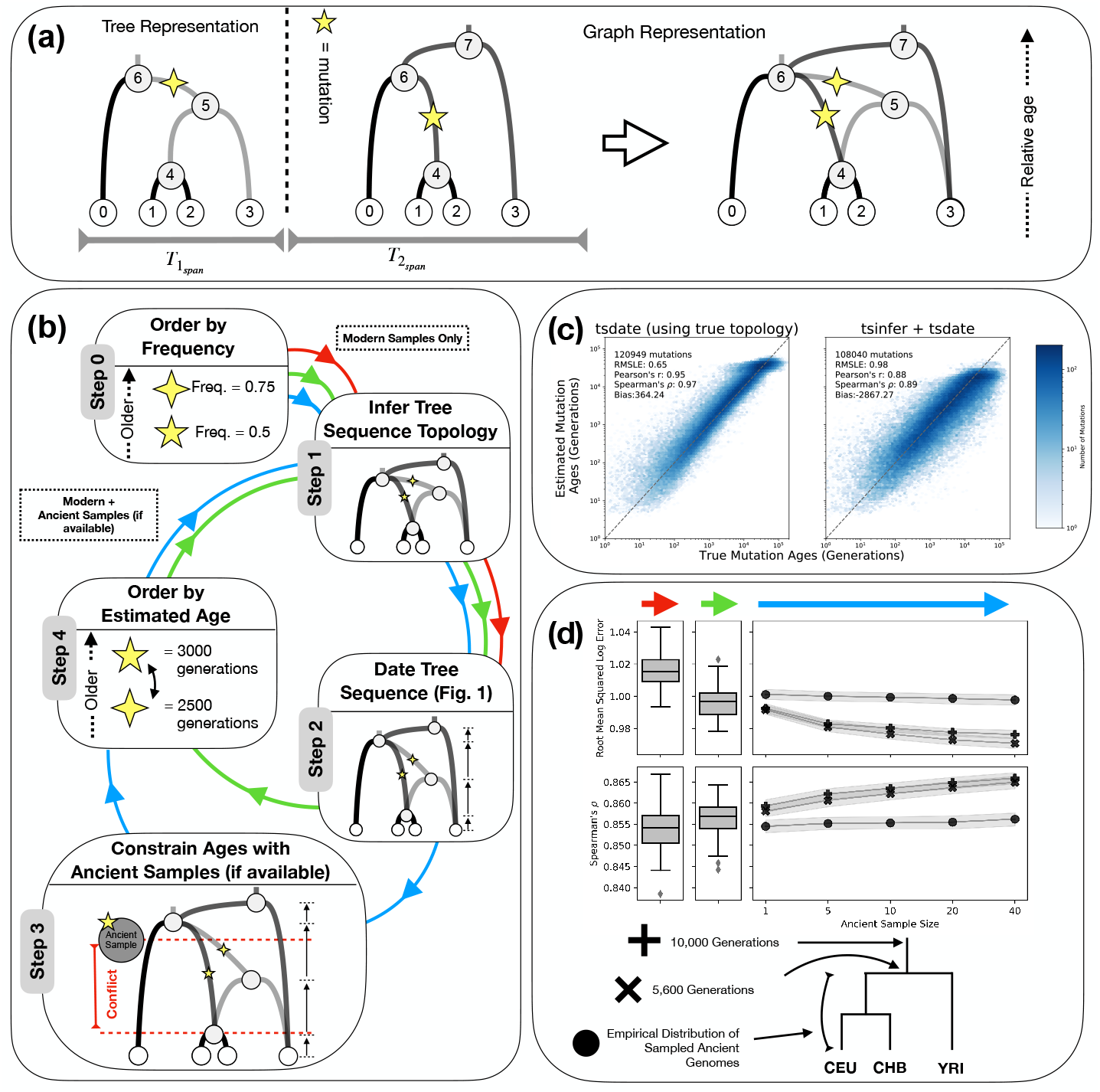
Schematic overview and validation of the inference methodology. (a) An example tree sequence topology with four samples (nodes 0-3), two marginal trees, and two mutations. *T*_1*span*_ and *T*_2*span*_ measure the genomic span of each marginal tree topology, with the dotted line indicating the location of a recombination event. The graph representation is equivalent to the tree representation. (b) Schematic representation of the inference methodology. Step 0: alleles are ordered by frequency; the mutation represented by the four-point star is thus considered to be older than that represented by the five-point star. Step 1: the tree sequence topology is inferred with `tsinfer` using modern samples. Step 2: the tree sequence is dated with `tsdate`. Step 3: node date estimates are constrained with the known age of ancient samples. Step 4: ancestral haplotypes are reordered by the estimated age of their focal mutation; the five pointed star mutation is now inferred to be older than the four-point star mutation. The algorithm returns to Step 1 to re-infer the tree sequence topology with ancient samples. Arrows refer to modes of operation: Steps 0, 1 and 2 only (red); after one round of iteration without ancient samples (green) and after one round of iteration with increasing numbers of ancient samples (blue). (c) Scatter plots and accuracy metrics comparing simulated (x-axis) and inferred (y-axis) mutation ages from neutral coalescent simulations from `msprime`, using `tsdate` with the simulated topology (left) and inferred topology (right); see Methods for details. (d) Accuracy metrics, root-mean squared log error (top) and Spearman rank correlation coefficient (bottom), with modern samples only (first panel), after one round of iteration (second panel) and with increasing numbers of ancient samples (coloured arrows as in panel b). Three classes of ancient samples are considered, reflecting where in history of humans they have been sampled from (see schematic below). See Supplementary Note S2.3 for details.

In this paper, we introduce, validate and apply non-parametric methods for inferring time-resolved tree sequences from multiple heterogeneous sources, building on previous work^10^ to efficiently infer a single, unified tree sequence of ancient and contemporary human genomes. We validate this structure using the known age and population affinities of ancient samples and demonstrate its power through spatio-temporal inference of human ancestry. We note that while humans are the focus of this study, the methods and approaches we introduce are valid for most recombining organisms.

### A unified genealogy of modern and ancient human genomes

To generate a unified genealogy of modern and ancient human genomes, we integrated data from eight sources. This included three modern datasets: the 1000 Genomes Project (TGP) which contains 2,504 sequenced individuals from 26 populations^1^, the Human Genome Diversity Project (HGDP), which consists of 929 sequenced individuals from 54 populations^9^, and the Simons Genome Diversity Project (SGDP) with 278 sequenced individuals from 142 populations^15^. 154 individuals appear in more than one of these datasets (see Supplementary Note S3). In addition, we included data from three high-coverage sequenced Neanderthal genomes^26–28^, the single Denisovan genome^29^, and newly reported high coverage whole genome data from a nuclear family of four (a mother, a father, and their two sons with average coverage of 10.8x, 25.8x, 21.2x, and 25.3x) from the Afanasievo Culture, who lived ∽ 4.6 thousand years ago (kya). Finally, we used 3,589 published ancient samples from over 100 publications compiled by the Reich Laboratory (see Supplementary Note S3) to constrain allele age estimates. These ancient genomes were not included in the final tree sequence due to the lack of reliable phasing for the majority of samples, though we later discuss a potential solution to this problem.

We first merged the modern datasets and inferred a tree sequence for each autosome using `tsinfer` version 0.2 (see Methods). This approach uses a reference panel of inferred ancestral haplotypes to impute missing data at the 95.6% of sites that have at least one missing genotype. We then estimated the age of ancestral haplotypes with tsdate (see Methods), a Bayesian approach that infers the age of ancestral haplotypes with accuracy comparable to or greater than alternative approaches and with unmatched scaling properties (see Fig. 1c, Extended Data Fig. 4-5). We identified 6,412,717 variants present in both ancient and modern samples. For each, a lower age bound is provided by the estimated archaeological date of the oldest ancient sample in which the derived allele is found. Where this was inconsistent with the initial inferred value (559,431, or 8.7% of variants) we used the archaeological date as the variant age.

Next, we integrated the Afanasievo family and four archaic sequences with the modern samples and re-inferred the tree sequences using the iterative approach outlined in Fig. 1b. In `tsinfer`, samples cannot directly descend from other samples (which is highly unlikely in reality). Instead, we create “proxy” ancestral haplotypes, which at non-singleton sites are identical to each ancient sample, and insert them at a slightly older time than the ancient sample. These proxies can act as ancestors of both modern and (younger) ancient samples. Thus, the ancient samples are never themselves direct ancestors of younger ones, but their haplotypes may be. The integrated tree sequences of each autosome together contain 26,958,720 ancestral haplotypes, 231,073,278 edges, 91,172,114 variable sites, and 245,631,834 mutations. We infer that 38.7% of variant sites require more than one change in allelic state in the tree sequence to explain the data. This may indicate either recurrent mutations or errors in sequencing, genotype calling, or phasing, all of which are represented by additional mutations in the tree sequence. If we discount mutations affecting only a single sample (indicative of sequencing error) we find that 13,513,873 sites contain at least two mutations affecting more than one sample, implying up to 17.5% of variable sites could be the result of more than one ancestral mutation. Fig. S3 shows that a high proportion of sites with over ∽ 100 mutations on chromosome 20 have sequencing or alignment quality issues as defined by the TGP accessibility mask^1^, or are in minimal linkage disequilibrium to their surrounding sites, suggesting they are largely erroneous. We chose to retain such sites to enable recovery of input data sources; however, future iterative approaches to the removal of such probable errors are likely to improve use cases such as imputation.

To characterise fine-scale patterns of relatedness between the 215 populations of the constituent datasets, we calculated the time to the most recent common ancestor (TMRCA) between pairs of haplotypes from these populations at the 122,637 distinct trees in the tree sequence of chromosome 20 (∽ 300 billion pairwise TMRCAs). After performing hierarchical clustering on the average pairwise TMRCA values, we find that samples do not cluster by data source (which would indicate major data artefacts), but reflect known patterns of global relatedness (Fig. 2 and Supplementary Interactive Fig. 1). We conclude that our method of integrating datasets is therefore robust to biases introduced by different datasets. Numerous features of human history are immediately apparent, such as the deep divergence of archaic and modern humans, the effects of the Out of Africa event (Fig. 2 (i)), and a subtle increase in Oceanian/Denisovan MRCA density from 2,000-5,000 generations ago (Fig. 2 (ii)). Multiple populations show recent within-group TMRCAs, suggestive of recent bottlenecks or consanguinity. The most extreme cases are for “populations” for which only a single individual exists in our dataset, such as the four archaic individuals, and a Samaritan individual from the SGDP. We find the Samaritan has an logarithmic average within-group TMRCA of ∽ 1,000 generations, which is caused by multiple MRCAs at very recent times (Fig. 2 (iii)) and is consistent with documented evidence of a severe bottleneck and extensive consanguinity in recent centuries^30^. Indigenous populations in the Americas, an Atayal individual from Taiwan, and Papuans also exhibit particularly recent within-group TMRCAs.

**Fig. 2:**
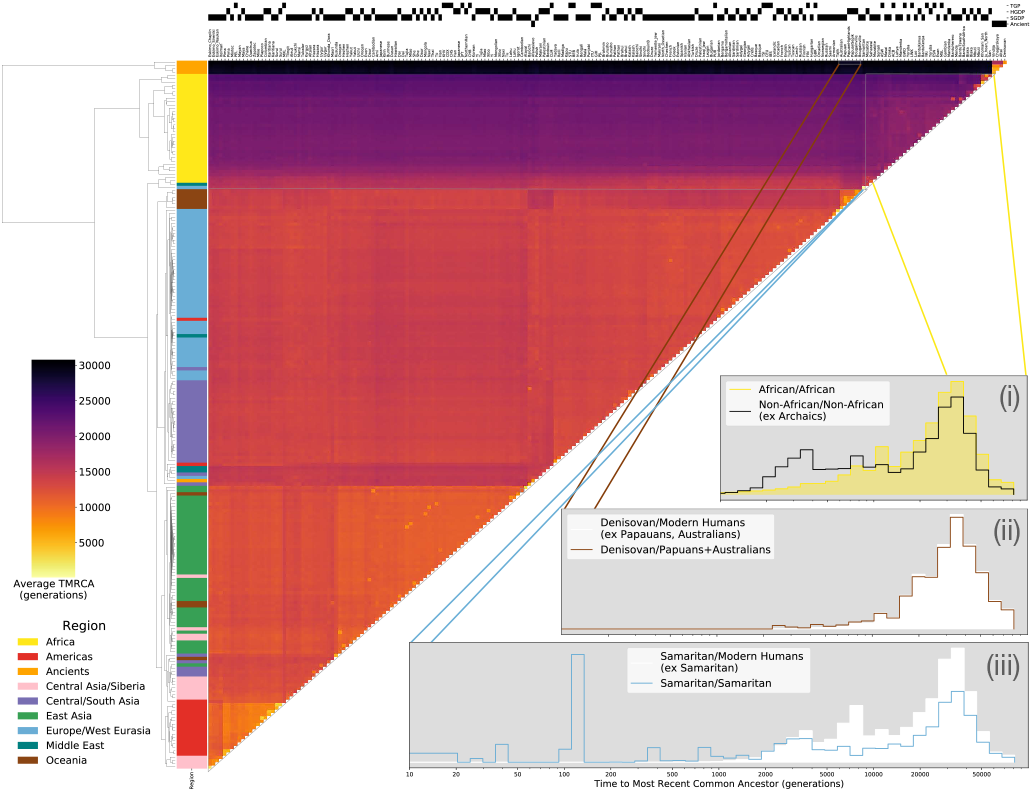
Clustered heatmap showing the average time to the most recent common ancestor (TMRCA) on chromosome 20 for haplotypes within pairs of the 215 populations in the HGDP, TGP, SGDP, and ancient samples. Each cell in the heatmap is coloured by the logarithmic mean TMRCA of samples from the two populations. Hierarchical clustering of rows and columns has been performed using the UPGMA algorithm on the value of the pairwise average TMRCAs. Row colours are given by the region of origin for each population, as shown in the legend. The source of genomic samples for each population is indicated in the shaded boxes above the column labels. Three population relationships are highlighted using span-weighted histograms of the TMRCA distributions: (i) average distribution of TMRCAs between all non-African populations (black line) compared to African/African TMRCAs (solid yellow). (ii) Denisovans and Papuan/Australian TMRCAs (solid line), compared to Denisovans against all non-Archaic populations (solid white). The subtle but unique signal is particularly evident in Supplementary Interactive Fig. 1 at https://awohns.github.io/unified_genealogy/interactive_figure.html). TMRCAs between the two Samaritan chromosomes (solid line), compared to the Samaritans/all other modern humans (solid white). Duplicate samples appearing in more than one modern dataset are included in this analysis.

### Tree-sequence based analysis of descent from ancient samples

To validate the dating methodology, we first used simulations to show that the integration of ancient samples improves derived allele age estimates under a range of demographic histories (Fig. 1d.). To provide empirical validation of the method, we considered the ability of the method and alternatives to infer allele ages that are consistent with observations from ancient samples. We inferred and dated a tree sequence of TGP chromosome 20 (without using the ancient samples) and compared the resulting point estimates and upper and lower bounds on allele age with results from `GEVA`^31^ and `Relate`^32^. This resulted in a set of 659,804 variant sites where all three methods provide an allele age estimate. Of these, 76,889 derived alleles are observed within the combined set of 3,734 ancient samples, thus putting a lower bound on allele age. We find that estimated allele ages from `tsdate` and `Relate` showed the greatest compatibility with ancient lower bounds, despite the fact that the mean age estimate from `tsdate` is more recent than that of `Relate` (Fig. 3a and Supplementary Note S5).

**Fig. 3:**
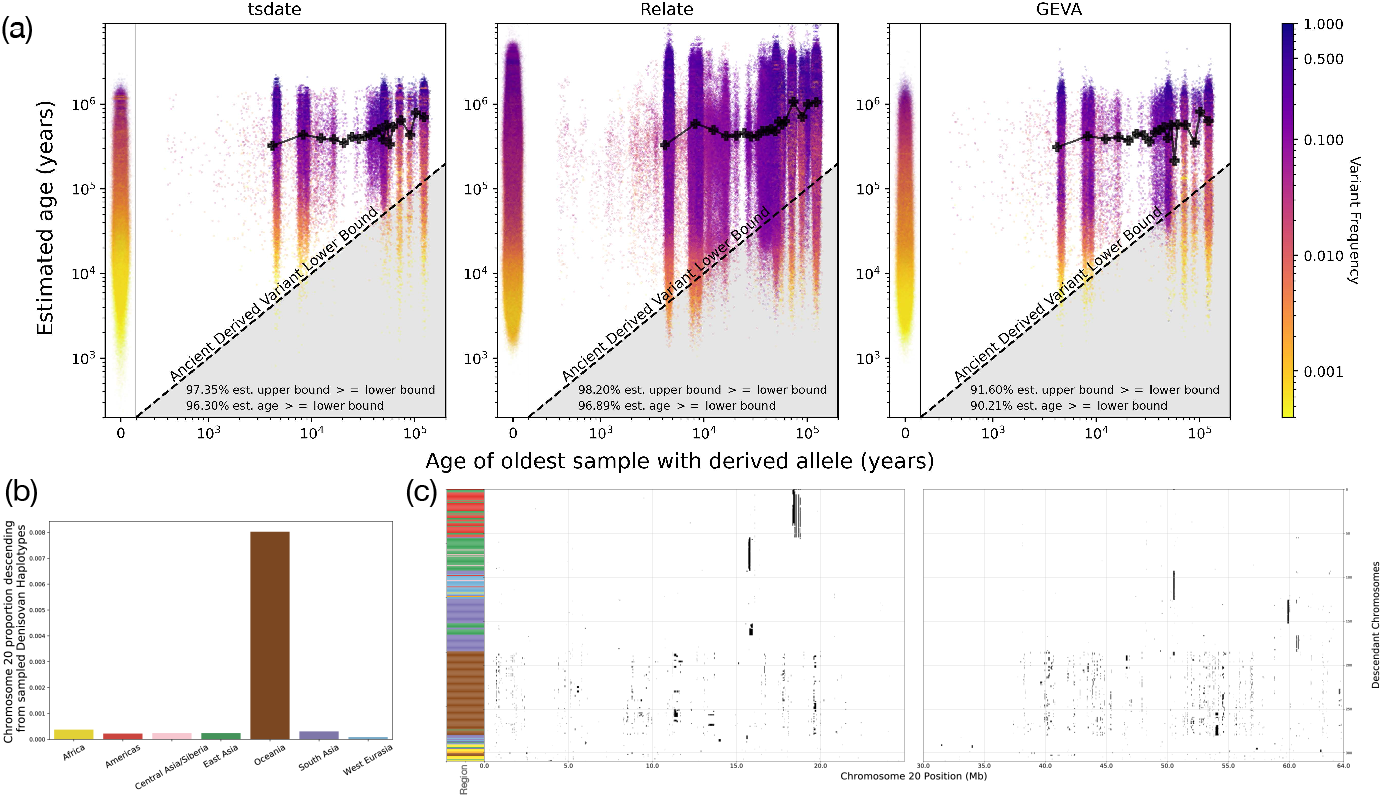
Validation of inference methods using ancient samples. (a) Comparison of mutation age estimates from three methods (`tsdate`, `Relate` and `GEVA`) using 3,734 ancient samples at 76,889 variants on chromosome 20. The radiocarbon-dated age of the oldest ancient sample carrying a derived allele at each variant site in the 1000 Genomes Project is used as the lower bound on the age of the mutation (diagonal lines). Mutations below this line have an estimated age that is inconsistent with the age of the ancient sample. Black lines on each plot show the moving average of allele age estimates from each method as a function of oldest ancient sample age. Plots to the left show the distribution of allele age estimates for modern-only variants from each respective method. Additional metrics are reported in each plot. (b) Percentage of chromosome 20 for modern samples in each region that is inferred to descend from “proxy ancestors” associated with the sampled Denisovan haplotypes, calculated using the genomic descent statistic^1^. (c) Tracts of descent along chromosome 20 descending from Denisovan “proxy ancestors” in modern samples with at least 100 kilobases (kb) of total descent (colour scheme as for Fig. 2).

Next, to assess the ability of the unified tree sequence to recover known relationships between ancient and modern populations, we considered the patterns of descent to modern samples from Archaic proxy ancestral haplotypes on chromosome 20. Simulations, detailed in Supplementary Note S2.6, indicate that this approach detects introgressed genetic material from Denisovans at a precision of ∽ 86% with a recall of ∽ 61%. We find that there are descendants among non-archaic individuals, including both modern individuals and the Afanasievo, for 13% of the span of the Denisovan proxy haplotypes on chromosome 20. The highest degree of descent among modern humans is in Oceanian populations as previously reported^29,33–35^ (Fig. 3b). However, the tree sequence also reveals how both the extent and nature of descent from the Denisovan proxy ancestors varies greatly among modern humans (Fig. 3c). In particular, we find that Papuans and Australians carry multiple fragments of Denisovan haplotypes that are largely unique to the individual. In contrast, other modern descendants of Denisovan proxy ancestors have fewer Denisovan haplotype blocks which are more widely shared, often between geographically distant individuals.

For the Afanasievo family, we find the greatest amount of descent from their proxy ancestral haplotypes among individuals in Western Eurasia and South Asia (Extended Data Fig. 6a), consistent with findings from the genetically similar Yamnaya peoples^36^. Notably, the most frequent descendant blocks in Extended Data Fig. 6b all contain geographically disparate modern samples. These cosmopolitan patterns of descent support a contemporaneous diffusion of Afanasievo-like genetic material via multiple routes^36^.

For the Neanderthal samples, where there are three samples of different age, our simulations indicate that interpretation of the descent statistics is complicated by varying levels of precision and recall among lineages. Nevertheless, recall is highest at regions where introgressing and sampled archaic lineages share more recent common ancestry and precision is higher for the most recent samples. Examining patterns of descent from the Vindija across autosomes indicates that modern non-African groups carry similar levels of Vindija-like material (see Extended Data Fig. 8), supporting suggestions that the proportions are very similar between East Asians and West Eurasians^37^ and inconsistent with other reports^27,38^.

### Non-parametric inference of spatio-temporal dynamics in human history

Tree-sequence based analysis of ancient samples demonstrates the power of the approach for characterising patterns of recent descent. To assess whether we could use the tree sequence to capture wider patterns in human history we developed a simple estimator of ancestor spatial location. We use the location of descendants of a node, combined with the structure of the tree sequence, to provide an estimate of ancestor location (see Methods). The approach can use information on the location of ancient samples, though it does not attempt to capture the geographical plausibility of different locations and routes. The inferred locations are thus a model-free estimate of ancestors’ location, informed by the tree sequence topology and geographic distribution of samples. Although the relationships between genealogies and spatial structure has been an active area of research in both phylogenetics and population genetics for many years^39–44^, our approach is the first to infer ancestral locations incorporating recombination. More sophisticated methods which also use genome-wide genealogies are currently in development, and show considerable promise (Osmond, M., and Coop, G., personal communication).

We applied the method to the unified tree sequence of chromosome 20, excluding TGP individuals (which lack precise location information). We find that the inferred ancestor location recovers multiple key events in human history (Fig. 4, Supplementary Video 1). Despite the fact that the geographic centre of gravity of all sampled individuals is in Central Asia, by 72 kya the average location of ancestral haplotypes is in Northeast Africa and remains there until the oldest common ancestors are reached. Indeed, the inferred geographic centre of gravity of the 100 oldest ancestral haplotypes (which have an average age of ∽ 2 million years) is located in Sudan at 19.4 N, 33.7 E. These findings reflect the depth of African lineages in the inferred tree sequence and are compatible with well-dated early modern human fossils from eastern and northern Africa^45,46^. We caution that our sampling of Africa is inhomogenous, and it is likely that if instead we analysed data from a grid sampling of populations in Africa the geographic centre of gravity of independent lineages at different time depths would shift. In addition, past major migrations such as the Bantu and Pastoral Neolithic expansions, both occurring within the last few thousand years, mean that present day distributions of groups in Africa and elsewhere may not represent ancestral ones, and thus the approach of using the present-day geographic distribution to provide insight is likely to give a distorted picture of ancient geographic distributions^47^. Nevertheless, this analysis demonstrates that the deep tree structure is geographically centred in Africa in autosomal data, just as it is for mitochondrial DNA and Y chromosomes^48,49^.

**Fig. 4:**
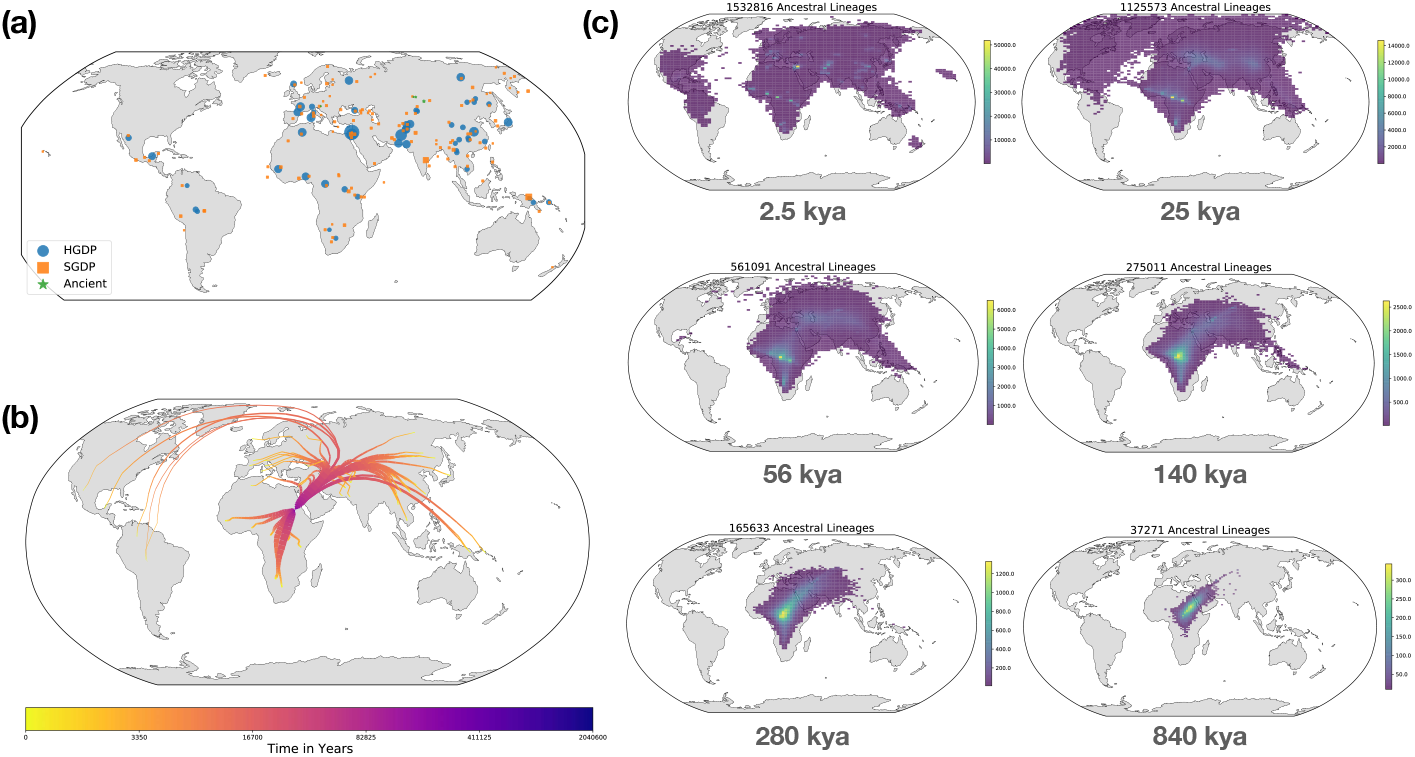
Visualisation of the non-parametric estimator of ancestor geographic location for HGDP, SGDP, Neanderthal, Denisovan, and Afanasievo samples on chromosome 20. (a) Geographic location of samples used to infer ancestral geography. The size of each symbol is proportional to the number of samples in that population. (b) The average location of the ancestors of each HGDP population from time t=0 to ∽ 2 million years ago. The width of lines is proportional to the number of ancestors of each population over time. The ancestor of a population is defined as an inferred ancestral haplotype with at least one descendant in that population. (c) 2d-histograms showing the inferred geographical location of HGDP ancestral lineages at six time-points. Histogram bins with fewer than 10 ancestors are not shown. Link to ancestral location video: https://www.youtube.com/watch?v=AvV0zBSdxsQ.

Traversing towards the present, by 280 kya, the centre of gravity of ancestors is still located in Africa, but many ancestors are observed in the Middle East and Central Asia and a few are located in Papua New Guinea. At 140 kya, more ancestors are found in Papua New Guinea. This is almost 100 kya before the earliest documented human habitation of the region^50^. However, our findings are potentially consistent with the proposed timescales of deeply diverged Denisovan lineages unique to Papuans^35^. At 56 kya, some ancestral lineages are observed in the Americas, much earlier than the estimated migration times to the Americas^51^. This effect is likely attributable to the presence of ancestors who predate the migration and did not live in the Americas, but whose descendants now exist solely in this region^52^; the same effect may also explain the ancient ancestors within Papua New Guinea. Additional ancient samples and more sophisticated inference approaches are required to distinguish between these hypotheses. Nevertheless, these results demonstrate the ability of inference methods applied to tree sequences to capture key features of human history in a manner that does not require complex parametric modelling.

## Discussion

A central theme in evolutionary biology is how best to represent and analyse genomic diversity in order to learn about the processes, forces and events that have shaped history. Historically, many modelling approaches focused on the temporal behaviour of individual mutation frequencies in idealised populations^53,54^. In the last 40 years there has been a shift toward modelling techniques that focus on the genealogical history of sampled genomes and that can capture the correlation structures in variation that arise in recombining genomes^24,55^. Critically, while allele frequency is an idealised and unknowable quantity, there does exist a single, albeit extremely complex, set of ancestral relationships that, coupled with how mutation events have altered genetic material through descent, describes what we observe today.

However, while the empirical generation of data has transformed our ability to characterise genomic variation and relatedness structures in humans, including that of ancient individuals, developing efficient methods for inferring the underlying genealogy has proved challenging^56,57^. Recent progress in this area^10,32^, on which we build, has been driven by using approximations that capture the essence of the problem but enable scaling to population-scale and genome-wide data sets. The methods described here produce high quality dated genealogies that include thousands of modern and ancient samples. These genealogies cannot be entirely accurate, nevertheless, they enable a wealth of novel analyses that reveal features of human evolution^25,58–60^. That our highly simplistic estimator of spatio-temporal dynamics of ancestors of modern samples captures key events, such as an East-African genesis of modern humans, introgression from now-extinct archaic populations in Asia and historical admixture^10^, suggests that more sophisticated approaches, coupled with the ongoing program of sequencing ancient samples, will continue to generate new insights into our history.

Moreover, because the tree sequence approach captures the structure of human relationships and genomic diversity, it provides a principled basis for combining data from multiple different sources, enabling tasks such as imputing missing data and identifying (and correcting) sporadic and systematic errors in the underlying data. Our results identified different types of error common in reference data sets (erroneous sites and genotyping error) as well as emphasising the importance of recurrent mutation in generating human genetic diversity^61,62^. Although additional work is required to correct such errors, as well as integrate other types of mutation, notably structural variation, a reference tree sequence for human variation - along with the tools to use it appropriately^10,25^ – potentially represents a basis for harmonising a much larger and wider set of genomic data sources and enabling cross data-source analyses. We note that such a reference tree-sequence could also enable data sharing and even privacy-preserving forms of genomic analysis^22^ through compression of cohorts against such a reference structure.

There exists much room for improvement in the methods introduced here, as well as new opportunities for genomic analyses that use the dated tree-sequence structure. For example, our approach requires phased genomes, which is a particular challenge for ancient samples that typically pick random reads to create a “pseudo-haploid genome”^63^. However, it should be possible to use a diploid version of the matching algorithm in `tsinfer` to jointly solve phasing and imputation. This also has the potential to alleviate biases introduced by using modern and genetically distant reference panels for ancient samples^64^. Recent work focusing on inferring genealogies for high-coverage ancient samples, and using mutations dated in such a genealogy to infer relationships of lower coverage samples through time, offers an alternative strategy for accommodating the unique challenges of ancient DNA in this context^65^. In addition, our approach to age inference within `tsdate` only provides an approximate solution to the cycles that are inherent in genealogical histories^66^ and there are many possible approaches for improving the sophistication of spatio-temporal ancestor inference.

## Methods

Novel algorithms are described in this section, while details of dataset preparation and simulations can be found in the Supplementary Notes.

### Tree sequence inference algorithm

`tsinfer` is a scalable method for inferring tree sequence topologies using genetic variation data^1^, which we update to version 0.2 by incorporating two features: provision for inexact matching in the copying process and support for missing data.

#### Mismatch, error, and recurrent mutation

The `tsinfer` algorithm is a two-step process. First, partial ancestral haplot-ypes for the sampled DNA sequences are constructed on the basis of shared, derived alleles at a set of sites. Second, a Hidden Markov Model (HMM) is employed from left to right along each haplotype to infer which, among the array of older haplotypes, gives the closest match. This is based on the Li and Stephens^2^ (LS) copying model. In previous versions of `tsinfer`, we supported only exact haplotype matching – if a haplotype matched perfectly against an ancestor up to a certain position, but then a mismatch occurred, it could only be explained by switching to a different ancestor via recombination. The new algorithm now supports the full LS model including a mismatch term which allows for inexact matching, where mismatches between a target haplotype and its inferred ancestor are explained by additional mutations. The relative probabilities of recombination versus mismatch are tuned via a “mismatch ratio” parameter: high ratios lead to fewer inferred recombination events and more additional mutations, while low ratios result in more recombination events and fewer additional mutations (see Supplementary Note S1.1).

Probabilities of recombination between adjacent sites can be provided in the form of a genetic map. However, optimal mismatch probabilities will depend on factors such as sequencing error and variation in mutation rates along the genome. To establish suitable mismatch ratios, we therefore evaluated inference performance on both simulated and real data. Extended Data Fig. 1 shows the effect of different mismatch probabilities, using a simulated 10 megabase (Mb) region of 1,500 human genomes, both with and without added error in sequencing and ancestral state polarisation. Different metrics disagree slightly on the optimal values used to minimise difference between the simulated and inferred trees (see Supplementary Note S1.2). However, error metrics are consistently low when mismatch ratios in both the ancestor and sample matching phases are set to between 0.001 and 10. This range is also suggested as optimal by two different proxy measures of tree sequence complexity based on file size. Extended Data Fig. 2 shows roughly the same pattern when inferring tree sequences from real data, although in these cases only file size measures are available ––– no ground truth exists for comparison. In all cases, good results are obtained by mismatch ratios close to unity, where the probability of a mismatch is set equal to the median probability of recombination between adjacent inference sites (marked as dashed and dotted lines on the plots). A mismatch ratio of 1 is therefore used in all further analyses; the breadth of the plateau in parameter space indicates similarly accurate results are obtained using mismatch ratios within an order of magnitude either side of this value.

#### Missing data

Missing data is accommodated in `tsinfer` by using older ancestors as a “reference panel” for imputation. In the core `tsinfer` HMM, samples and ancestors copy from older ancestors. If the sample or ancestor contains missing data at a site, the missing genotypes are imputed from the most recent ancestor without missing data. The approach provides a principled approach to imputing missing data for both contemporary and ancient genomes, as only older ancestral haplotypes are used.

### Dating algorithm

We use the tree sequence topology estimated by `tsinfer` as the basis for an approximate Bayesian method to infer node age. This approach is implemented in the open-source software package `tsdate`.

#### Conditional coalescent prior

The first step in the algorithm is to assign a prior distribution to the age of ancestral nodes in the tree sequence. A coalescent prior is an obvious choice^3–5^. However, rather than use a fully tree sequence-aware prior, we use an approximate approach based on assigning marginal priors to each node. Specifically, we use the number of modern samples descending from each node to find a mean age and associated variance under the coalescent^6^. In the tree sequence, a single node may span many trees, and therefore be associated with several of these means and variances: we take the average, weighted by tree span, resulting in an average mean and average variance for each ancestral node. We then use moment matching to fit a lognormal distribution as a prior, *π_u_*, for the age of node *u* in the tree sequence. Details of this approach can be found in Supplementary Note S1.2.

#### Time discretisation

Our inference approach requires a time grid for efficient computation. This is constructed by taking the union of the quantiles of the prior distribution of each ancestral node. The advantage of this approach is that inference is focused on times with greater probability under the prior, outperforming a naive, uniform grid. The density of the time grid is determined by the user-specified number of quantiles to draw from each ancestral node as well as a value, *ε*, which establishes the minimum time distance between points in the grid. The conditional coalescent prior *π_u_* for a node *u* allows us to find a probability *π_u_*(*t*) for each time-slice *t* in the grid.

### Inside-Outside algorithm

With a prior in place for ancestral nodes in the tree sequence and a time grid, we infer the age of nodes using a belief propagation approach we call the inside-outside algorithm, based on an HMM where the age of nodes are hidden states. In the case of a single tree, this equates to the standard forward-backward algorithm. In the case of a tree sequence, we must also consider the relative genomic spans associated with edges and deal with cycles in the undirected graph underlying the tree sequence. Cycles occur whenever a node has multiple parents and present a general problem in belief propagation^7^.

The algorithm is efficient because it uses dynamic programming and the tree sequence traversal methods implemented in `tskit`, the tree sequence toolkit. Scaling is linear with the number of edges in the tree sequence (Extended Data Fig. 5) and quadratic with the number of time slices used.

#### Inside pass

We seek to compute all values in the inside matrix *I* for all nodes and times in the discretised time grid. *I_u_*(*t*) is the probability of node *u* at time *t*, which encompasses the probability of all nodes and edges in the subgraph beneath *u*.

We initialise the prior probability of a sample node to be 1 at its sampled time and 0 elsewhere. We then proceed backwards in time, using the relationships between nodes encoded in the `tskit` edge table until we reach the most recent common ancestor (MRCA) nodes of the tree sequence. For each node, we visit every child as well as every time *t* in the time grid using

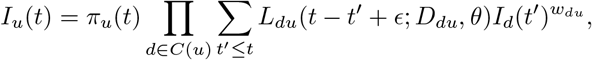

where *C*(*u*) is the set of all child nodes of *u*, and as previously defined, *π_u_*(*t*) is the prior probability of node *u* at time *t*. *L_du_*(*t − t′* + *ε*; *D_du_, θ*) is the mutation-based likelihood function of the edge from focal node *u* at time *t* to child node *d* at time *t′*. *ϵ* is an arbitrarily small value that is used to prevent parent and child nodes from existing at the same time slice. *D_du_* is the data associated with the edge including the span of the edge and the number of mutations on it. *θ* is the population-scaled mutation rate. *w_du_* is the span of the edge leading from *u* to *d* divided by *s_d_*, the total span of node *d* in the tree sequence. Note that the inside probability of node *d* is geometrically scaled by *w_du_* to address overcounting if *d* has multiple parents.

The likelihood function gives the probability of observing *k* mutations on an edge of length *δt* = *t − t′* + *ϵ* with span *l_du_*. It is Poisson distributed with parameter (*θl_du_δt*)*/*2

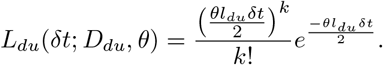

We can factorise the inside probability as

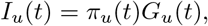

where *G_u_*(*t*) is

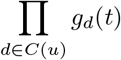

and *g_d_*(*t*) is

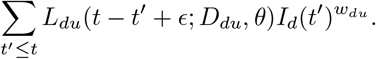

This factorisation will be useful in describing the outside pass in the next section. The equation terminates at the MRCA(s) of the tree sequence. The total likelihood of the tree sequence is obtained by taking the product of the inside matrix of each MRCA.

#### Outside pass

Once we have iterated up the tree sequence to find the inside matrix at every node, the inside probability of the MRCAs contain all of the information encoded in the tree sequence. To find the full posterior on node age, we now take account of the information in the tree sequence “outside” of the subgraph of each ancestral node. While this algorithm empirically performs well with a single inside and outside pass, any cycles in the underlying undirected graph (which occur when recombination causes a node to have more than one parent) will result in overcounting. The alternative “outside-maximisation” pass introduced in Supplementary Note S1.2.2 provides another approximate solution in these cases, though we find that the outside pass performs better empirically (see Fig. S2).

Beginning with the MRCA nodes in each marginal tree (the roots), we initialise the outside value of these nodes, *O_MRCAs_*, to be one at all non-zero time points. There is no information “outside” the MRCAs because all information in the tree has already been propagated to the node and is encoded in the MRCAs’ inside matrices. In a tree sequence, it is possible for a node *u* to be the MRCA in some of the marginal trees in which it appears but not in other trees. In these cases we find *O_u_* by dividing the span of trees where the node is the MRCA by *s_u_*, the total span of *u* in the tree sequence.

We then proceed down the tree sequence (forwards in time), again using the edge table sorted in descending order by the children’s age. At every node we calculate:

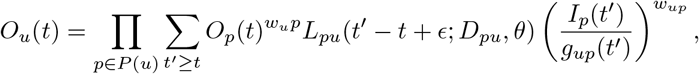

where *P*(*u*) is the set of parents of node *u* and other terms are defined in the previous section on the inside algorithm.

Once the inside and outside passes are complete, the approximate posterior can be calculated as

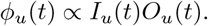

Importantly, the mean value of the posterior distribution may not be consistent with the tree sequence topology. We provide the option to “constrain” node age estimates by forcing each node to be older than the estimated age of its children. The unconstrained mean and variance of each node are retained as metadata in the tree sequence. The full posterior can also be retained separately if desired.

We observe that mutations mapping to edges descending from the single oldest root in tree sequences inferred by `tsinfer` are generally of lower quality, so in our implementation of the outside pass we include an option to avoid traversing such edges. We use this setting in all analyses using `tsdate` in this work. Additionally, results from tsdate do not include estimates for mutations appearing on these edges.

### Iterative approach for inferring tree sequences with ancient and modern samples

We combined `tsinfer` and `tsdate` in an iterative approach that allows for the incorporation of ancient samples and improves inference accuracy in many settings.

The first step of the iterative approach is to order derived alleles appearing in the sample by their frequency. `tsinfer` requires a relative ordering of derived alleles to both build ancestral haplotypes and infer copying paths. Frequency is a largely accurate and highly efficient means of providing an ordering for these ancestors^1^. Once alleles are ordered, it is possible to infer a tree sequence topology with `tsinfer` (Fig. 1b Step 1).

With an inferred tree sequence topology, we next estimate the age of inferred ancestral haplotypes with `tsdate` (Fig. 1b Step 2). If using tsdate’s outside pass, we do not constrain the resulting date estimates by the topology.

If ancient samples are present, we can use them to constrain the estimated age of derived alleles. The previous step (Fig. 1b Step 2) provides date estimates for the inferred ancestors as well as for mutations. Since we estimate the age of the ancestral nodes above and below a mutation, the child node of an edge hosting a mutation is constrained by the ancient sample-informed lower bound on derived allele age. This bound is determined by gathering the haplotypes of ancient samples (either sequenced or genotyped) and examining derived alleles that can be called in these ancient samples with high confidence. If multiple ancient samples carry the same derived allele, we use the oldest sample as the lower bound on its age. Once lower bounds have been collected for all derived alleles observed in ancient samples, we compare these with our statistically inferred lower bounds on allele age, adjusting our age estimates where necessary to ensure consistency with ancient samples. Any radiocarbon-dated ancient samples with high-confidence variant calls may provide constraints in this step, including unphased and/or low-coverage samples. Although a substantial fraction of radiocarbon dates are likely inaccurate, we note that there is a low probability that errors on the order of a few thousand years will meaningfully affect tree sequences inferred using this approach. Only a subset of erroneously dated alleles will be older than the true age of the mutation, which would affect allele age estimation accuracy, and still fewer will be older than the ancestral haplotype from which they descend, which would affect topological estimation accuracy.

With allele age estimates from step 2, possibly constrained by ancient samples in step 3, we are now able to re-infer the tree sequence topology. The revised age estimates are used to order the age of derived alleles when re-estimating ancestral haplotypes with `tsinfer`; if they are more accurate than frequency in determining a relative ordering of mutations, topological inference accuracy should be improved. Indeed, we find that the iterative approach improves accuracy when re-inferring tree sequences from variation data simulated with a uniform recombination map and without error (Fig. 1d). When reinferring tree sequences from data simulated with error or with a variable recombination map, less improvement is observed (Fig. S4).

Ancient samples can be included in tree sequences that are (re)-inferred with estimated allele ages. This is accomplished by inserting ancient samples at their correct relative ordering among ancestors generated by `tsinfer`. Only phased ancient samples with an age estimate may be included, although we note that extending `tsinfer`’s HMM to handle diploid individuals may allow for phasing of ancient samples in this step. We additionally produce “proxy ancestors” associated with ancient samples at a slightly older time than the ancient sample. These are composed of all non-singleton sites carried by the ancient sample, and may serve as ancestors to younger ancestors and samples. Finally, we infer copying paths between ancestors and samples to produce a tree sequence of modern and ancient samples.

### Inferring the Location of Ancestors in a Tree Sequence

We use a naive, non-parametric approach to gain insight into the geographic location of ancestral haplotypes based on the known locations of sampled genomes. The latitude and longitude coordinates of individual samples are provided for SGDP individuals, while the location of sampling centres were used for the HGDP individuals. No geographic information was provided for TGP individuals, so these were not used in location inference. We also used the coordinates of the archaeological sites associated with the Afanasievo and Archaic individuals.

The weighted centre of gravity is determined for each ancestral node by iterating up the tree sequence, visiting child nodes before their parents using the same traversal pattern as for the previously described inside algorithm. At each focal ancestral node, we find the geographic midpoint between each of the children of that node. The following simple approach was used to find the geographic midpoints. For a node *u*, the latitude and longitude coordinates of the child node of each edge descending from *u* were converted to Cartesian coordinates. We find the average of the children’s coordinates and convert this back to latitude and longitude. We then continue up the tree sequence using this location to calculate the coordinates of *u*’s parents.

This method is highly efficient, requiring less than 1 minute to compute on the combined tree sequence of chromosome 20.

## Data Availability

Newly reported sequencing data from the Afanasievo family is available from the European Nucleotide Archive, accession number PRJEB43093; phased variant data for the family can be downloaded from https://reichdata.hms.harvard.edu/pub/datasets/release/wohn_2021_phasedAfanasievo/. All publicly available datasets used in this paper are available from their original publications. See Supplementary Note for details.

## Code Availability

`tsinfer` is available at https://tsinfer.readthedocs.io/ under the GNU General Public License v3.0, tsdate at https://tsdate.readthedocs.io/ under the MIT License, and tskit at https://tskit.readthedocs.io/ under the MIT License. All code used to perform analyses in this paper can be found at https://github.com/awohns/unified_genealogy_paper.

## Acknowledgements

Funded by the Wellcome Trust (grant 100956/Z/13/Z to GM), the Li Ka Shing Foundation (to GM), the Robertson Foundation (to JK), the Rhodes Trust (to AWW), the NIH (NIGMS grant GM100233 to DR), the Paul Allen Foundation (to DR), the John Templeton Foundation (grant 61220 to DR) and the Howard Hughes Medical Institute (to DR). The computational aspects of this research were supported by the Wellcome Trust (Core Award 203141/Z/16/Z) and the NIHR Oxford BRC. The views expressed are those of the authors and not necessarily those of the NHS, the NIHR or the Department of Health. We thank the Oxford Big Data research computing team, specifically Adam Huffman and Robert Esnouf, and Daniel Lieberman and E. Castedo Ellerman for comments.

## Author Contributions

We used the CRediT contributor roles taxonomy (https://casrai.org/credit/): A.W.W.: Conceptualization, Data curation, Formal analysis, Investigation, Methodology, Project administration, Resources, Software, Validation, Visualization, Writing—original draft, Writing—review & editing. Y.W.: Conceptualization, Data curation, Formal analysis, Investigation, Methodology, Resources, Software, Supervision, Validation, Visualization, Writing—original draft, Writing—review & editing. B.J.: Data curation, Formal analysis, Investigation, Validation, Visualization. A.K.: Data curation, Formal analysis, Methodology, Resources. S.M.: Data curation, Formal analysis, Resources. R.P.: Investigation, Methodology, Resources. N.P.: Methodology. D.R.: Funding acquisition, Investigation, Project administration, Resources, Supervision, Writing—review & editing. J.K.: Conceptualization, Funding acquisition, Methodology, Resources, Software, Supervision, Writing—original draft, Writing—review & editing. G.M.: Conceptualization, Funding acquisition, Methodology, Supervision, Writing—original draft, Writing—review & editing.

## Competing Interests

GM is a director of and shareholder in Genomics plc and a partner in Peptide Groove LLP.

---

**Extended Data Fig. 1:**
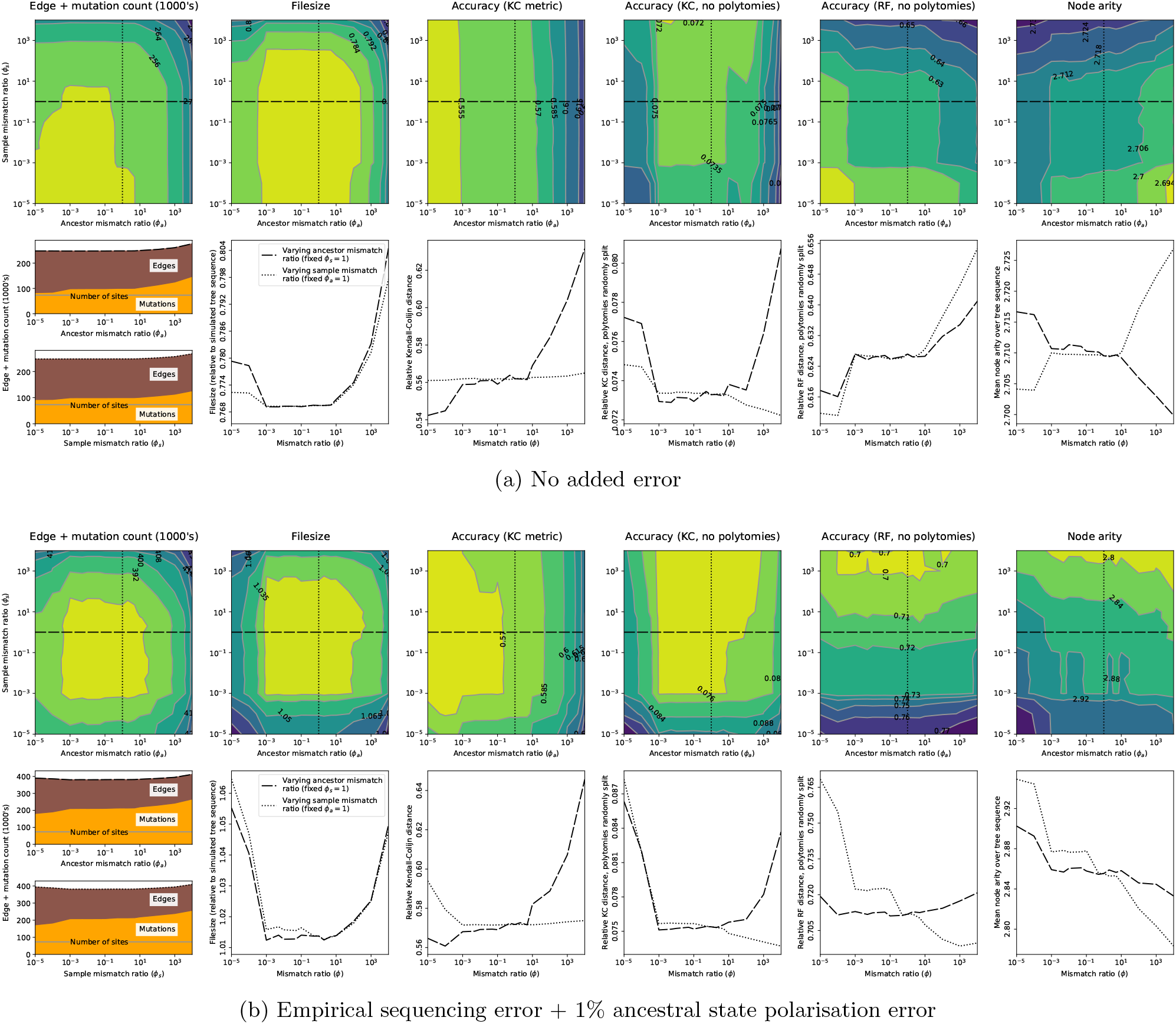
The effect of varying the mismatch ratio on accuracy metrics for `tsinfer`. Results from 1,500 simulated human-like genome sequences of 10 Mb in length. (a) Simulations without error. (b) Simulations with an empirically calibrated genotyping error model and 1% error in ancestral state assignment. For each panel, the upper (coloured contour) plots show accuracy metrics as a function of the mismatch ratio in ancestor matching (x-axis) and in sample matching (y-axis) algorithms. Slices through contour plots indicated by the dashed and dotted lines are plotted in the lower (line) plots. The total number of edges plus mutations, and filesize relative to the simulated tree sequence (first 2 columns) are indirect measures of accuracy. Direct measures of inference accuracy provided via the Kendall-Colijn (KC) or Robinson-Foulds (RF) tree-distance metrics (middle columns) which can, however, be influenced by polytomy size (i.e. node arity: last column); breaking polytomies at random may reduce this influence. Metrics are normalised against maximum expected distances. See Supplementary Note S2.1 for further details.

**Extended Data Fig. 2:**
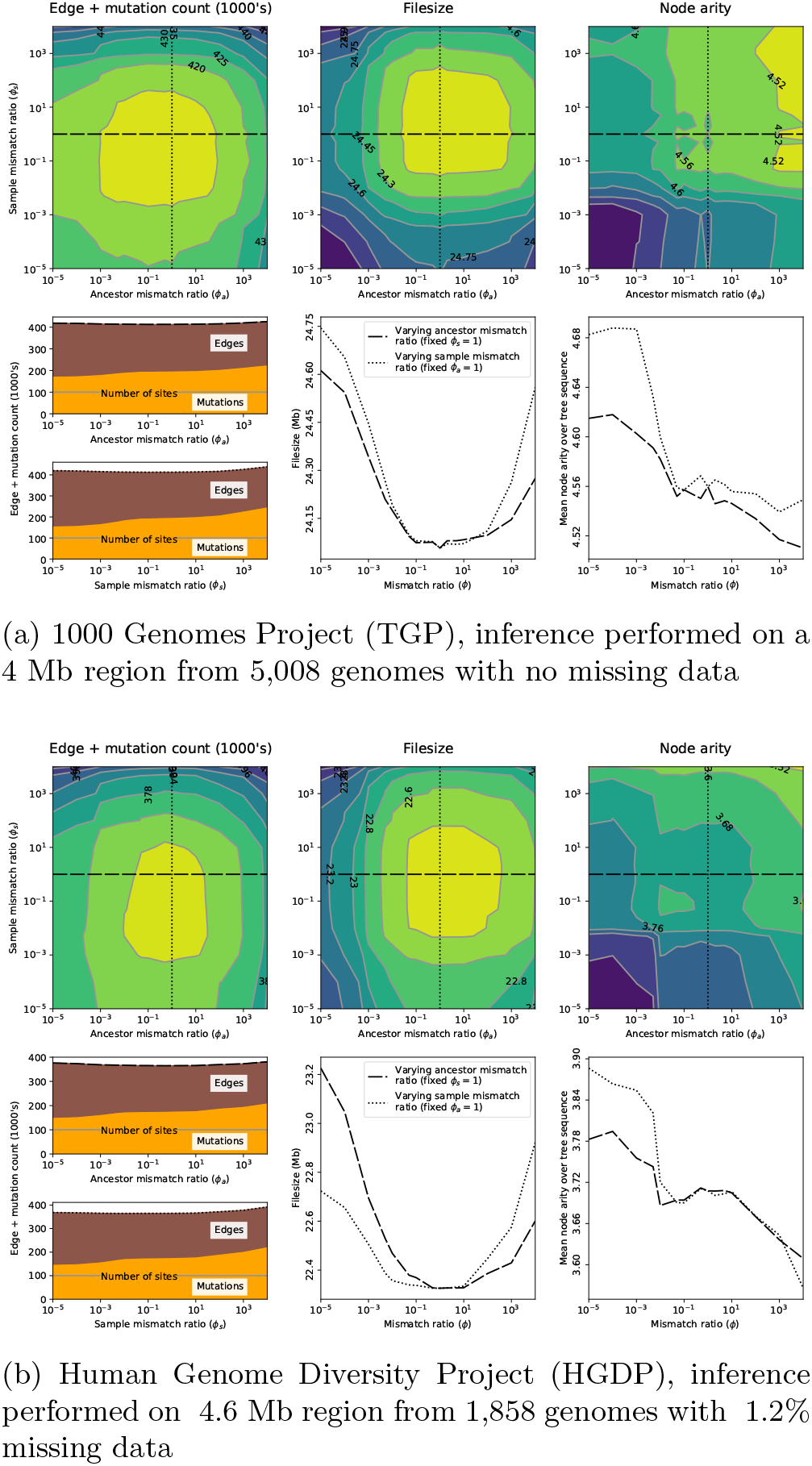
Effect of mismatch ratio parameter on `tsinfer` results from empirical sequence data (based on a subset of 100,000 sites on the short arm of chromosome 20). Dotted and dashed lines as for Extended Data Fig. 1. See Supplementary Note S2.1 for further details.

**Extended Data Fig. 3:**
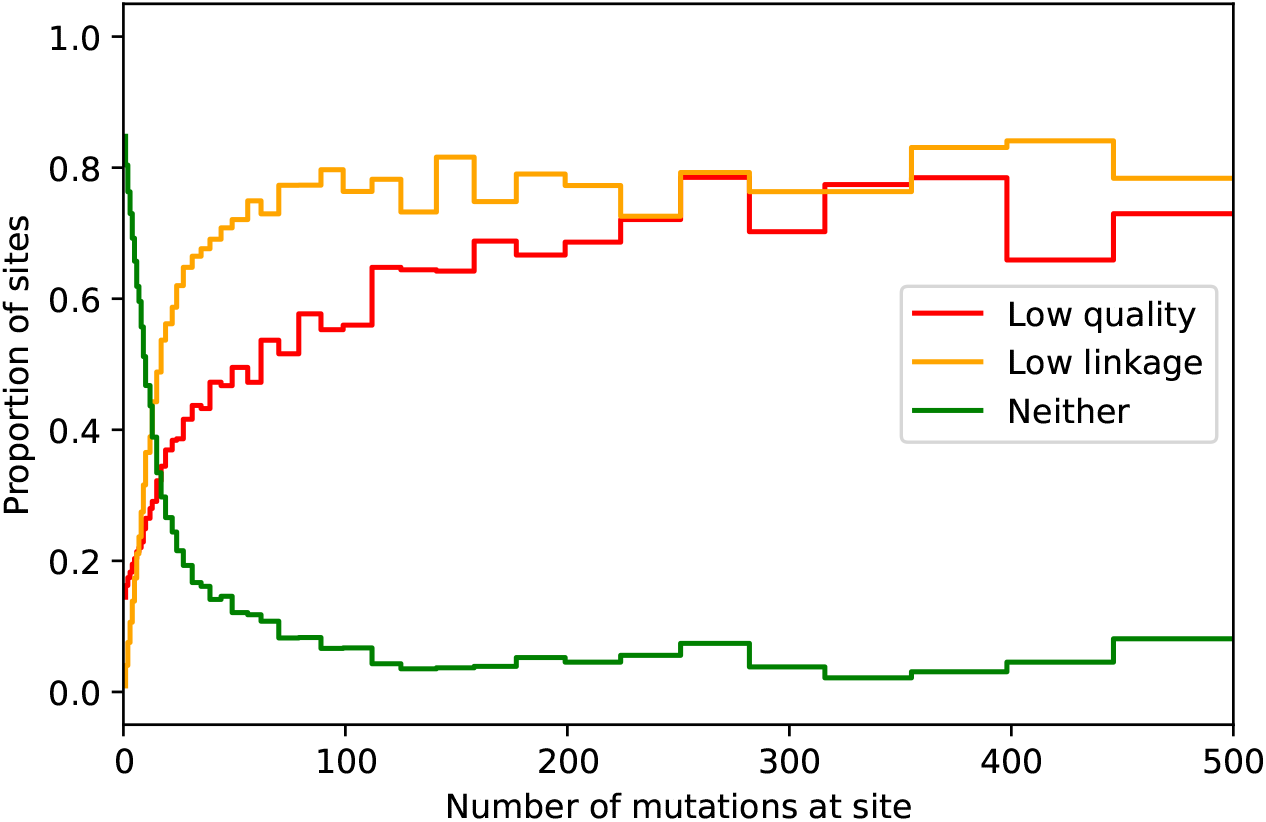
The relationship between metrics of variant-calling error and the number of mutations on the inferred tree sequence. For the combined tree sequence of HGDP, TGP, SGDP, and ancient samples, the proportion of sites in two categories - low quality and low linkage, binned by the number of mutations at the site. Low quality is defined by the TGP strict accessibility mask. Sites that fail any of the accessibility mask filters such as low coverage or low mapping quality are marked as low quality. Low linkage is defined by summing the linkage disequilibrium (*r*^2^) for the 50 sites either side of the focal site. If this quantity is less than 10 the site is marked as low linkage.

**Extended Data Fig. 4:**
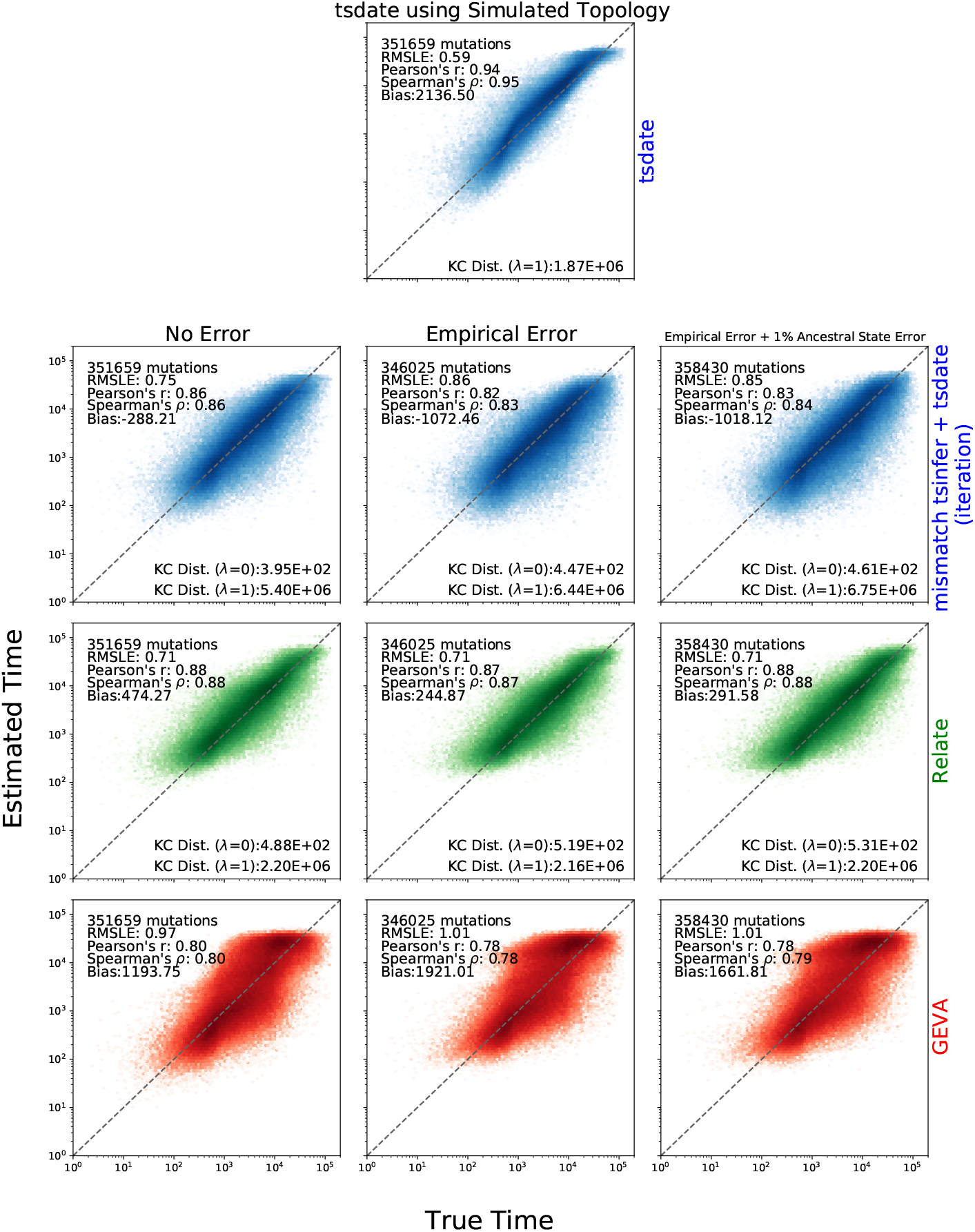
Accuracy of variant age inference. Evaluation of the accuracy of `tsdate`, `tsinfer`, `GEVA` and `Relate` on 5 Mb sections of chromosome 20 simulated using the `OutOfAfrica_3G09` model implemented in `stdpopsim`^2–4^ with the chromosome 20 GRCh37 recombination map and 100 samples each from YRI, CEU, and CHB. 30 replicates were performed. The top row shows the accuracy of `tsdate` on the simulated topology. The second row shows the results of inferring tree sequences with `tsinfer`, dating the tree sequence with `tsdate`, and then re-inferring and re-dating the tree sequence (using the chromosome 20 recombination map and a mismatch ratio of 1). The third row shows the results of `Relate` using a script provided by the authors to re-infer branch lengths and Ne. The fourth row shows the results of `GEVA` with default parameters. The first column shows simulations without genotype error, the second shows simulations with an empirical error model and the third shows an empirical error model and 1% ancestral state error. In each column, only sites dated by all three methods are shown. Summary metrics are shown in each subplot.

**Extended Data Fig. 5:**
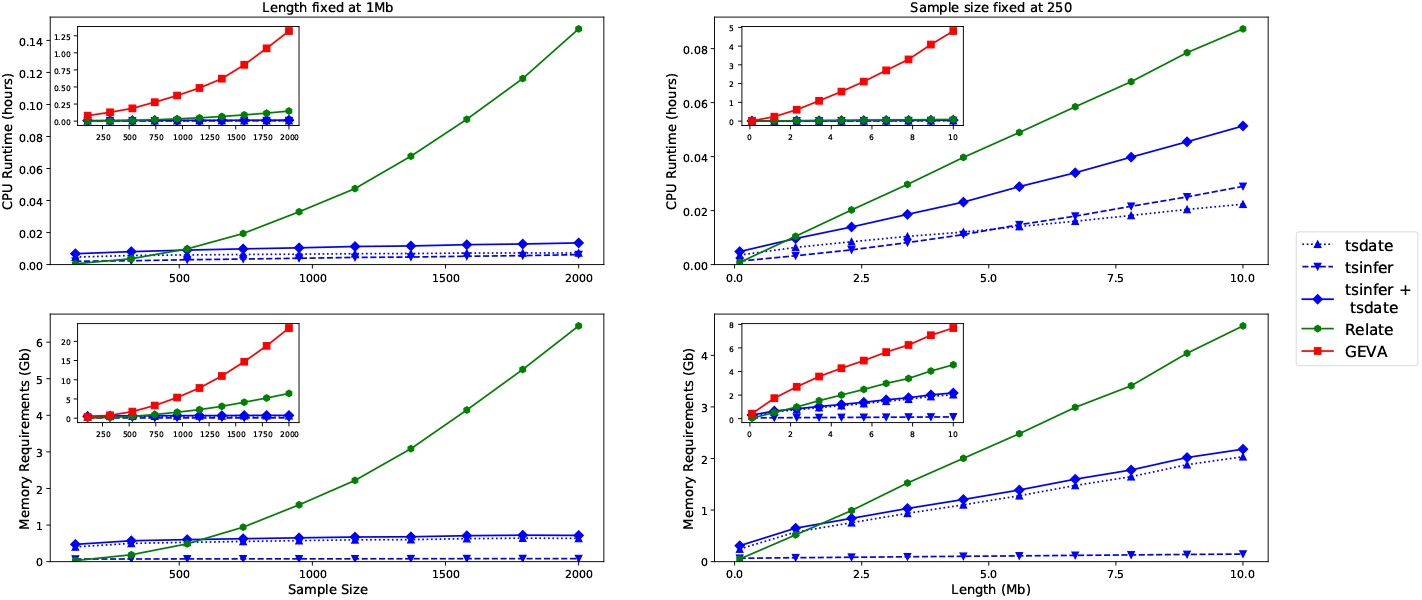
Scaling properties of `tsdate` compared to `tsinfer`, `Relate`, and `GEVA`. The left column shows the CPU and memory requirements for inference using the three methods on `msprime` simulations of 1 Mb, with *N_e_* = 10^4^, *μ* = *r* = 10^*−*8^ and sample sizes from 10 to 2000 (`Relate` continues to scale quadratically with larger sample sizes). Five replicates were performed at each sample size. The column on the right shows results of ten `msprime` simulations with sample size fixed at 250 and simulated lengths of 100 kb to 10 Mb. The main axes compare results for `tsdate`, `tsinfer`, and `Relate`. The inset plots show the same data with the addition of `GEVA`, which otherwise obscures the differences between other methods.

**Extended Data Fig. 6:**
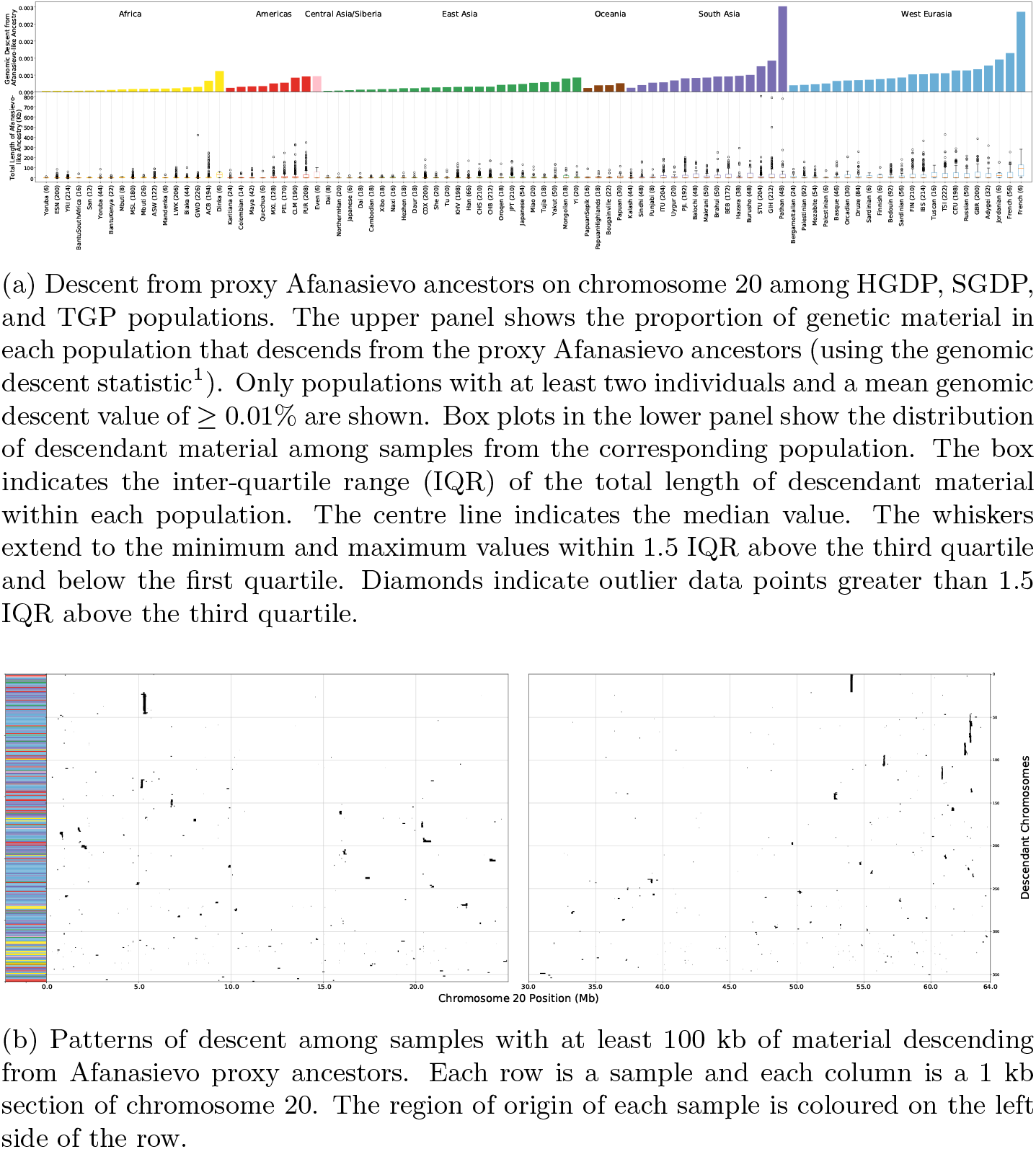
Inferred patterns of descent from Afanasievo proxy ancestors on chromosome 20. The colour scheme for each region is the same as in Fig. 2.

**Extended Data Fig. 7:**
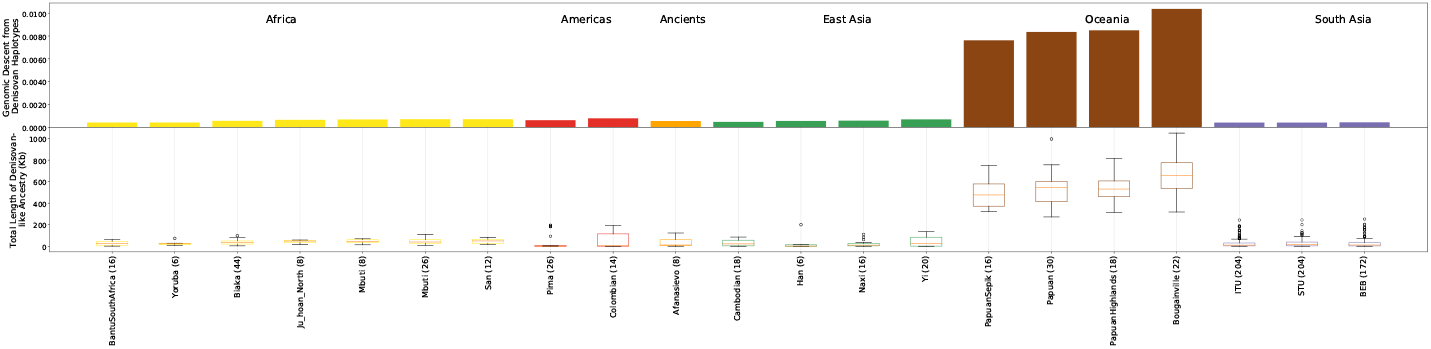
Descent from Denisovan proxy ancestors on chromosome 20. The upper portion shows the proportion of genetic material in each population that descends from the proxy Denisovan ancestors (the genomic descent statistic^1^). Box plots in the lower panel show the distribution of descendant material among samples from the corresponding population; box plot elements are defined in Fig. 6a. Only populations with a mean genomic descent value of ≥ 0.04% are shown. See Fig. 3 for details of the modern haplotypes inherited from Denisovan proxy ancestors.

**Extended Data Fig. 8:**
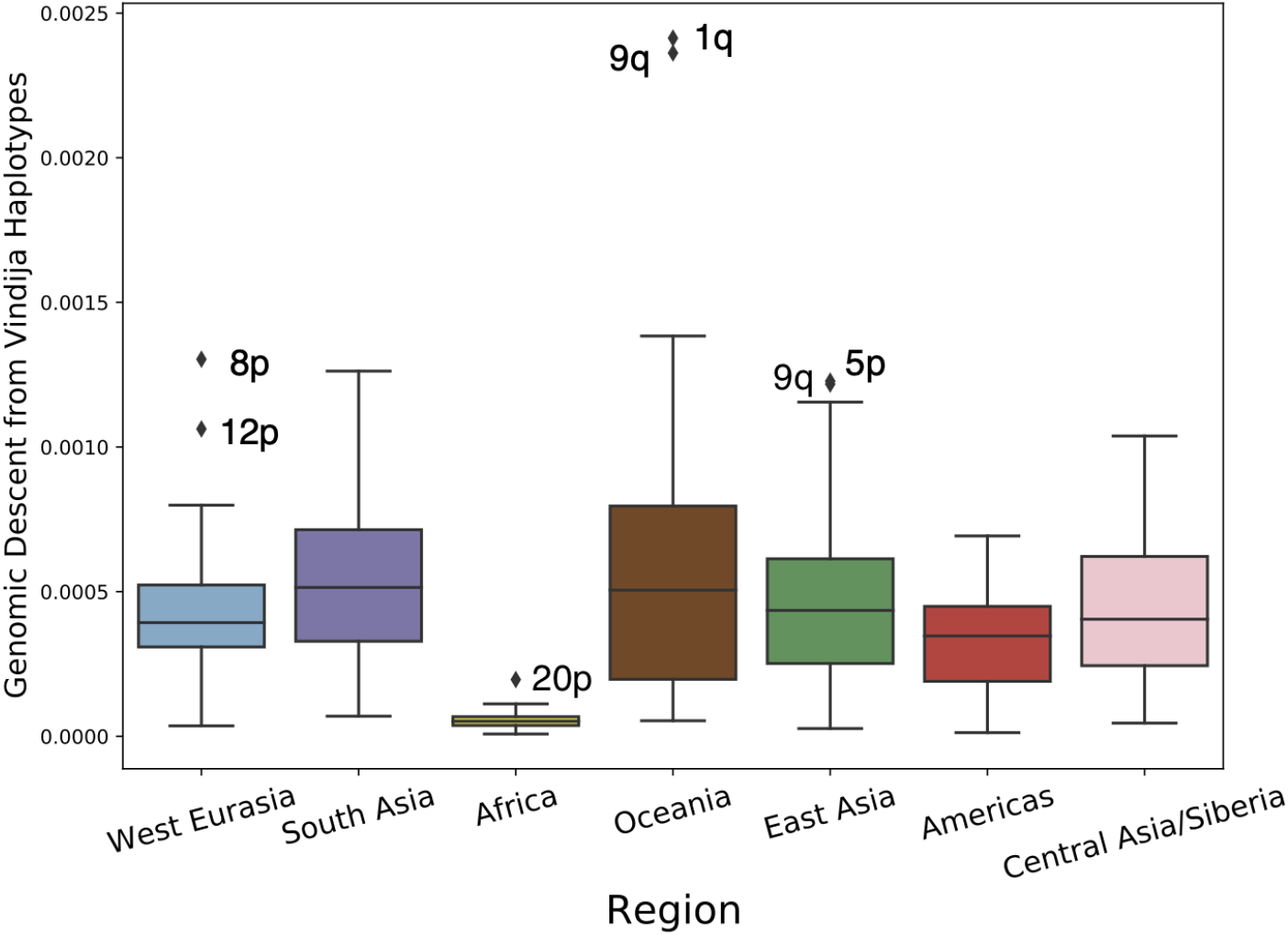
Descent from Vindija Neanderthal proxy haplotypes. Each data point is a genomic descent statistic^1^ calculated from the inferred tree sequence of the arms of each autosome. The statistic gives the proportion of genetic material in modern samples from each region that descends from Vindija proxy haplotypes. Box plot elements are defined in Fig. 6a. Outliers are labelled.

## Notes

https://awohns.github.io/unified_genealogy/interactive_figure.html

https://www.youtube.com/watch?v=AvV0zBSdxsQ&feature=youtu.be

