## Supplementary Note for "A unified genealogy of modern and ancient genomes"

### S1 Methods Detail

#### S1.1 Inferring tree sequence topology using `tsinfer` with mismatches

The `tsinfer` matching algorithm is based on a Hidden Markov Model (HMM) in which a hidden state is inferred at an array of positions ("inference sites") along a haplotype[1]. The hidden state is the identity of the most recent ancestor of the current haplotype, from among the $n$ possible ancestors at that position. In other words, the hidden state specifies the ancestor of which the haplotype is an immediate copy. Finding the most likely array of hidden states — the most likely "copying path" for a haplotype — requires us to specify HMM *transition* and *emission* probabilities; these correspond to the effects of recombination and mismatch (arising from error or mutation) respectively.

Transition probabilities are relevant when moving from one position to the next along the genome. With a recombination rate (i.e. genetic distance, or mean number of crossovers) of $r$ between the current and previous position, as calculated from a standard genetic map[2], the probability, $p_r$, of a detectable recombination is given by the standard map function[3]

$$p_r = (1 - e^{-2r})/2. \tag{1}$$

As in the Li and Stephens formulation[4,5], if a recombination occurs, the hidden state may switch to any of the $n$ ancestors with equal probability, giving transition probabilities of $p_r/n$ to any alternative state, and a probability of $1 - p_r + p_r/n$ of the state remaining unchanged[6].

From `tsinfer` version 0.2, emission probabilities are included in the HMM matching algorithm. These allow for mismatch between the hidden state and the observed haplotype: an emission probability of 0 means that the observed allele at a site is always that indicated by the hidden state, an emission probability of 1 means the observed allele never matches that indicated by the hidden state. For biallelic sites with $a = 2$ alleles, an emission probability of ½ thus indicates that the identity of the ancestor at this site has no influence on the observed allele. As it is only the *relative* values of emission probabilities, compared to the transition probabilities, that are important, we parameterize emission via a *mismatch ratio*, $\phi$. This is used to calculate a single emission probability used at all sites, $m$, as a function of the median recombination rate between adjacent inference sites, $\tilde{r}$. More specifically, we set

$$m = (1 - e^{-a\phi\tilde{\rho}})/a. \tag{2}$$

For biallelic sites ($a = 2$) and small $\tilde{r}$, Equation 2 returns an emission probability approximately $\phi$ times the median recombination probability ($\tilde{p_r}$). If a mismatch is inferred from the Viterbi decoding of the forward algorithm, it is incorporated into the resulting tree sequence by placing an additional mutation in the haplotype being matched. Note, however, that such mutations may either reflect true recurrent or back mutations, or represent errors of various sorts, such as in sequencing or in the approximations made in the `tsinfer` algorithm.

#### S1.2 `tsdate`

The `tsdate` algorithm is an approximate Bayesian method for inferring the age of ancestral nodes in a tree sequence topology. There are two main components to the algorithm: the construction of a prior on node age and propagation of likelihoods up and down the tree sequence using the inside-outside algorithm. The latter is described in detail in the methods. Here, we explain the prior in greater detail as well as provide an alternative to the outside pass: the outside-maximisation pass.

##### S1.2.1 Conditional Coalescent Prior

The coalescent provides a natural prior for the age of nodes in a tree sequence. However, the number of lineages remaining at any given time is unknown, so we instead use the theoretical results of Wiuf & Donnelly (1999) to find the mean and variance of node age under the "conditional coalescent." This involves conditioning on the number of samples descending from each ancestral node.

We begin by considering a coalescent tree with $n$ sequences, partitioned into $i$ and $n - i$. We condition on the event $E$ that all samples in $i$ coalesce before sharing a common ancestor with a sample outside of $i$. Conditioning on $E$ affects the order of coalescence events, but not the exponentially distributed waiting times. We seek to find the mean and variance of the time $\tau$ when all $i$ samples have coalesced. At this time there will be only one ancestor of $i$ remaining, but depending on how many lineages outside of $i$ have coalesced, the number of total ancestors, $\alpha$, could range from 2 to $n-i+1$.

Equation (10) in Wiuf & Donnelly (1999) shows that the expectation of $\tau$ is

$$E(\tau) = \frac{i-1}{n}. \tag{3}$$

The variance can be found using Equation 3 and the expectation of $\tau^2$, which is calculated as

$$E(\tau^2) = \sum_{m=2}^{n-i+1} P(\alpha = m)E(\tau^2|\alpha = m),$$

where $E(\tau^2|\alpha)$ is given by Equation 10 in Wiuf & Donnelly (1999) as

$$E(\tau^2|\alpha) = 8\sum_{j=\alpha}^{n} \frac{1}{j^2} + \frac{8}{n} - \frac{8}{\alpha} - \frac{8}{n\alpha},$$

and $P(\alpha = m)$ is Corollary 2 of Wiuf & Donnelly (1999) when only one ancestor in $i$ remains:

$$P(\alpha = m) = \frac{\binom{n-m-1}{i-2}\binom{m}{2}}{\binom{n}{i+1}}.$$

Coalescent simulations using `msprime` show that the distribution of the logarithm of the age of a node with a fixed number of descendant samples is approximately normal. We can thus use the method of moments to fit a lognormal distribution to the mean and variance of the age of each variant under the coalescent:

$$\sigma = \sqrt{log\left(\frac{Var(\tau)}{E(\tau)^2} + 1\right)};$$

$$\mu = log(E(\tau)) - \frac{1}{2}\sigma^2.$$

Although the prior is valid at any given marginal tree topology in the tree sequence, recombination introduces two complications: nodes may not span the full tree sequence and a given node may have descendant subsamples of differing sizes in the various marginal trees in which it appears. To address these points, we first iterate through each marginal tree in the tree sequence to find the total length of sequence *spanned* by node $u$, which we will refer to as $s_u$. We then determine the "sample weights" for each node by finding the span of sequence where $u$ has a particular number of descendent samples as a proportion of $s_u$. Since neighbouring marginal trees in a tree sequence are correlated, we use the difference in edges between each adjacent tree as a highly efficient method for calculating the sample weights.

Once we have evaluated sample weights for each node in the tree sequence, we construct a mixture distribution for the prior on node age. This is accomplished by finding the weighted average of expectation and variance of the prior distribution using the sample weights. With the weighted means and variances, we again use the method of moments to determine the parameters of the matched lognormal distribution.

We show that the lognormal approximation is well calibrated to data simulated under the coalescent both with and without recombination in Fig. S1.

##### S1.2.2 Outside-Maximisation Pass

Cycles in the undirected graph underlying a tree sequence occur when a node has more than one parent as a consequence of recombination (an example is shown in Fig. 1a). These cycles result in the over-counting of information in the outside pass of the inside-outside algorithm. To account for this, we introduce the outside-maximisation pass as an alternative. The rationale for this approach is that the inside value of the MRCA(s) of the sample has accumulated all of the data encoded in the tree sequence. In our model, the age of a node and the age of all non-descendant nodes are conditionally independent, given the age of a node's parents. Thus, if we fix the age of the MRCA(s), we can then walk down the tree sequence fixing the age of parent nodes before considering the age of their children.

First, we set the age of each MRCA in the tree sequence to the time slice with the maximum probability in the node's inside matrix (equivalent to assigning the age with the greatest maximum posterior probability). Next, we establish a traversal path using the topology such that each node is only visited once all of its direct and indirect parents have been considered. For each node we visit on this traversal path, we know the age of all of a node's ancestors as well as the relative likelihood of the subtree below each node for a given age. For any possible node age in the discretised time grid we can also compute the likelihood of the events on the branches to its direct and already-dated ancestors, as described in the inside pass. With this information we can compute the conditional posterior density for the age of the node. We set its age to the time slice with the maximum value.

Formally, for each node $u$ we seek to calculate the time of the node, $T_u$ as

$$T_u = \underset{t \leq U_p}{\arg\max} \left( I_u(t) \prod_{p \in P(u)} L_{pu}(T_p - t + \epsilon; D_{pu}, \theta) \right),$$

where $U_P$ is the age of the youngest parent of node $u$ and other variables are defined in the Methods section on the inside and outside passes.

#### S2 Evaluation of Methods

Code implementing all evaluations can be found at `https://github.com/awohns/unified_genealogy_paper`.

#### S2.1 Evaluating the accuracy of topological inference

There are two phases of matching in `tsinfer`: firstly matching inferred ancestors against older ancestors, secondly matching the known sample haplotypes against all ancestors[1]. Two mismatch ratios can therefore be specified, one in the ancestor-matching phase ($\phi_a$) and the other in the sample-matching phase ($\phi_s$). We expect optimal ratios to depend on the degree to which assumptions in the inference algorithm are met (for example, the assumption that frequency of the derived allele can be used as a proxy for ancestral age), and the amount of error in the analysed dataset (for example, a suitable value of $\phi_s$ should make inference reasonably robust to sequencing error). To find appropriate mismatch ratios, we performed inference for a range of 15 mismatch ratios from $10^{-5}$ to $10^{4}$ in both ancestor- and sample-matching phases, using a variety of datasets. Results are shown in Extended Data Figs. 1 (simulated datasets) and 2 (real datasets).

Simulated sequences were obtained from a standard three population "Out of Africa" model[7], as implemented in `stdpopsim`[8] version 0.1.2, with uniform mutation and recombination rates of $1.29 \times 10^{-8}$ /bp/gen and $1.72 \times 10^{-8}$ /bp/gen respectively. 500 chromosomes of 10 megabases (Mb) in size were sampled from each of the populations in the model, for a total sample size of $n = 1,500$. Accuracy of inference can be directly assessed by comparing the topology of the inferred trees along the genome with the ground-truth topologies. However, inferred trees contain polytomies (nodes whose number of direct children, i.e. arity, is greater than two), which have different effects on different tree distance metrics. For this reason, we compare accuracy using three separate metrics: the Kendall-Colijn (KC) metric[9] with $\lambda = 0$, the same metric but with each inferred tree having polytomies resolved equiprobably into a randomly chosen bifurcating topology, and finally the Robinson-Foulds (RF) metric[10]. The two KC metrics are normalised by dividing by the value obtained when comparing the ground truth tree with a "star" topology in the first case, and a random binary topology in the second. The RF metric, which is notoriously sensitive to minor tree changes but which has been shown to perform reasonably in practice[11], is normalized against the maximum possible number of disagreeing splits for two bifurcating trees ($2n - 4$): for this reason the RF with randomly split polytomies was used, although the metric without polytomy splitting gives similar qualitative results (data not shown). We can also assess inference performance indirectly by looking at file size or numbers of edges and mutations, under the assumption that lower values provide a more parsimonious representation of evolutionary history. This is particularly important when assessing performance on real data, for which ground-truth topologies are not available.

As expected, for simulations with no injected error, the first KC plot and the RF plot indicate that the lower the mismatch ratio, the greater the accuracy (Extended Data Fig. 1a); the KC metric suggests this effect is stronger for $\phi_a$, whereas the RF metric suggests it is (slightly) stronger for $\phi_s$. In contrast, where inferred trees have had their polytomies split at random, the KC metric suggests that very low mismatch ratios, particularly in the ancestor matching phase, are suboptimal: this is also suggested from the size of the resulting tree sequence. This difference may be due to the effect of polytomy size (arity of nodes), as the KC metric returns lower values when comparing a resolved tree with a polytomy than the average value comparing the same tree with the polytomy split equiprobably into binary resolutions. Also as expected, as mismatch ratios tend to zero, the number of mutations tends to the minimum possible (i.e. the number of sites): however there is a notable decrease in the number of mutations (and concomitant increase in the number of edges) when mismatch ratios drop from $10^{-3}$ to $10^{-4}$; this is also associated with an increase in file size.

In seeking optimal mismatch ratios for general use, it is more relevant to consider inference in which simulated sequences have had error added. Extended Data Fig. 1b gives results in which the sequences produced by simulation have had error added before inference. Genotyping error was added on the basis of estimates from the platinum genomes project[12] and ancestral states were flipped at a randomly chosen 1% of variant sites. As expected, with these injected errors, better results were obtained with higher mismatch ratios. The RF metric suggests an optimal $\phi_a$ between $10^{-4}$ and 10 but a much higher $\phi_s$. The KC metric with polytomies retained suggests lower $\phi_a$ values and $\phi_s$ somewhere between $10^{-3}$ and $10^2$. However, both the file size and the count of edges plus mutations clearly indicate an optimal range between $10^{-3}$ and 10 for both $\phi_a$ and $\phi_s$, a range that is mirrored in the case of $\phi_a$ by the KC metric with randomly resolved polytomies. From these results we deduce that none of the tree distance measures that we use are an ideal measure of inference accuracy, but that all agree that a mismatch ratio in the ancestor matching phase between $10^{-3}$ and 10 provides good inference. Moreover, within this range, the exact value is of minor importance. In the sample matching phase, the RF and plain KC metrics conflict over whether mismatch ratios higher than 10 are of benefit. However, such high values are not only contraindicated by measures of file size, but also because we would not expect error rates to be orders of magnitude higher than recombination rates. The simulated data therefore leads us to conclude that the value 1 is a reasonable mismatch ratio in both the ancestor and sample matching phases, that both file size and number of mutations plus edges are sensible proxies for inference accuracy, and that similar results would be obtained by using mismatch ratios anywhere between 0.1 and 10.

The proposed mismatch ratios of $\phi_a = \phi_s = 1$ are also supported by inference of real sequence data from two public datasets: the Thousand Genomes Project data (using build 37 of the human genome reference sequence), and the Human Genome Diversity Project (using build 38). The appropriate genetic map for chromosome 20 was used to specify transition probabilities via Equation 1, and for reasons of computational efficiency, inference was performed after subsetting down to sites 1,000,000 to 1,100,000 . Extended Data Fig. 2 shows that mismatch ratios between 0.1 and 10 provide the smallest file sizes and most parsimonious number of mutations and edges, with the newer and less error-prone HGDP dataset able to retain small sizes for slightly smaller values of $phi_s$. Reassuringly, the pattern in node arity reflects that in Extended Data

Fig. 1b, with larger polytomies for low values of $\phi_a$ and $\phi_s$. This indicates that, for the `tsinfer` algorithm, our method of injecting error into simulated datasets produces a similar effect to error in real data.

Optimal mismatch ratios around 1 might initially appear to be unreasonably high: recombination is expected to be common, recurrent mutations rare, and in modern datasets error should affect only a small fraction of the genotype matrix. However, simulations show that reasonable results are obtained even for mismatch ratios well above 1, where mismatches are substantially more probable than recombination events. This indicates that a correctly inferred recombination event can remove the need for several mismatches at nearby sites.

#### S2.2 Evaluating the accuracy of the `tsdate` prior

We evaluate the accuracy of the `tsdate` coalescent prior in Fig. S1. As described in Section S1.2.1, `tsdate` uses results from the coalescent conditioned on the frequency of an ancestral haplotype[13] to determine the mean and variance of the age of each node in the tree sequence. Moment matching is then used to fit a lognormal distribution on the age of each node. We evaluate the accuracy of this prior distribution, and also an alternative using a gamma distribution approximation, in simulations with and without recombination. The results of `msprime` simulations under the neutral coalescent show that the 95% credible interval of the gamma prior includes the true node ages more frequently than the equivalent lognormal distribution. However, the credible interval of the gamma is much wider than the lognormal and skews towards younger ages. For this reason, the lognormal is the default setting in `tsdate`.

#### S2.3 Accuracy of Age Estimation

Simulation-based evaluation of the accuracy and scaling properties of `tsinfer` and `tsdate` were performed with `msprime` version 1.0,[14] and `stdpopsim`[8]. We evaluate neutral coalescent simulations with recombination as well as a more complex model based on the Out of Africa event[7] (also used in Section S2.1). The effects of genotype and ancestral state errors are modelled using an error model based on an empirical comparison of the Illumina Platinum Genomes Project to matched data from the TGP[12]. Our methods are compared to `Relate`[15], a leading method for inferring dated genealogies and `GEVA`[12], a method for estimating allele age based on pairwise haplotype comparisons.

`tsdate` was evaluated under a variety of inference settings. First, the "true" topologies produced by `msprime` were passed to `tsdate`. This is indicated in relevant figures as "`tsdate` (using simulated topology)". The second method of evaluation was to run `tsdate` on tree sequence topologies inferred from the simulated variation data with `tsinfer`. This is referred to as "`tsdate` using `tsinfer` topologies". Both the outside and outside-maximisation passes are assessed in Fig. S2, as are various ratios of mutation to recombination rates. Since the outside pass empirically outperforms the outside-maximisation pass, only results using the former are used in subsequent simulations as well as in all results using real data. Variation data was converted to the required input formats for `GEVA` and `Relate` before running these programs using default settings. Mutation rates of $10^{-8}$ per base pair per generation are passed to all inference methods and recombination maps are passed to `tsinfer` and `Relate`. Ground truth allele ages were provided by `msprime`. For `tsdate` and `Relate`, allele age point estimates were determined using the arithmetic mean of the ages of the upper and lower bounding ancestors. The `tsdate`-estimated age of these ancestors was given by the mean of each node's posterior distribution. The mean age of the joint clock with quality-filtered pairs was used for `GEVA`. Only sites that could be dated by all methods were evaluated. This means that singletons, $n-1$ tons, and other sites not dated by one or more methods were excluded from the evaluation.

Root mean squared log error (RMSLE), Pearson's r, Spearman's $\rho$, and estimator bias were used to assess the accuracy of allele age estimates. Tree inference accuracy was measured by comparing simulated and inferred tree sequences using the KC tree distance metric[9], which includes a parameter, $\lambda$, determining the relative weight given to tree topology vs branch lengths when performing comparisons. We test both $\lambda = 0$, where all weight is given to the topology, and $\lambda = 1$, where all weight is given to branch lengths. See section S2.1 for details on the advantages and disadvantages of this metric.

Fig. S3 shows that our methods infer genealogies and allele ages with accuracy comparable or superior to the state-of-the-art in neutral coalescent simulations. When using the simulated topologies, `tsdate` has high accuracy in recovering allele age. When using inferred topologies from `tsinfer`, the accuracy of allele age estimation is comparable to `GEVA` and `Relate`.

Fig. S4 shows the performance of `tsdate` on simulations of chromosome 20 using the Out of Africa model[7,8] and the previously described error models. Demography impacts allele age estimates since `tsdate` uses a constant value of $N_e$ and does not currently model population structure. Using mismatch terms in `tsinfer` results in improved topological inference in all simulations and improvements in allele age accuracy by most metrics. The iterative approach shows variable effects depending on the simulation model used and evaluation metric being considered. Inference using the iterative approach provides the lowest values of KC distance with $\lambda = 1$ (corresponding to the highest accuracy in branch length estimation); however, KC distance with $\lambda = 0$ and the various allele age accuracy measures show that the iterative approach generally only offers comparable or slightly improved accuracy compared to a single pass of `tsinfer` with mismatch terms. However, the simulations in Section S2.5 show that when inferring genealogies from data simulated with a uniform recombination map, iteration improves accuracy. Inference using error-prone variation data leads to lower accuracy by most measures. These same patterns are observed in Extended Data Fig. 4, which compares the accuracy of our methods with `GEVA` and a version of `Relate` that re-estimates effective population size and branch lengths.

#### S2.4 Scaling Simulations

We evaluated the scaling properties of `tsinfer` and `tsdate` compared to `Relate` and `GEVA` using `msprime` neutral coalescent simulations; Extended Data Fig. 5. The CPU run-time and memory requirements are recorded using two evaluation schemes. First, the length of the simulated chromosomes is held constant while increasing sample sizes are used. Second, sample size is fixed while sequence length increases. Default parameters are used for all methods. The maximum memory allowance for `Relate` was set to 32 Gb to allow the scaling analysis to proceed without error.

#### S2.5 Ancient Samples and Allele Age Estimation Accuracy

Fig. 1d shows a simulation-based evaluation of allele age inference accuracy when incorporating ancient samples into the previously described Out of Africa model[7,8]. We simulated 5 Mb of chromosome 20 with a uniform recombination map and sampled 1008 modern samples split evenly between the three populations of the Out of Africa model: Yorubans (YRI), Han Chinese (CHB), and Utah Residents with Northern and Western European Ancestry (CEU). 40 ancient samples were also sampled. Three ancient sampling schemes were evaluated: (1) pick samples at times randomly selected from the empirical distribution of the ages of published ancient samples, (2) pick samples from the human population immediately prior to the time of the Out of Africa event in this model (5,650 generations ago), and (3) pick samples from 10,000 generations ago. 20 simulation replicates were performed with each sampling scheme.

The first sampling scheme used the estimated ages of ancient samples from the Reich Laboratory's compiled dataset of published ancient samples available at `https://reich.hms.harvard.edu/downloadable-genotypes-present-day-and-ancient-dna-data-compiled-published-papers`. The ancient sample times ranged from 90 years to 90,000, with a mean of 4,553 years (standard deviation: 4,554 years). If the randomly sampled ages were younger than 21,200 years (the split time of CEU and CHB in the Out of Africa model), they were assigned to CEU or CHB with 50% probability. If older than the split time, they were assigned to the $N_B$ population as described in Gutenkunst et al. (2009). All mutations only carried by ancient samples were removed from the simulation.

We then inferred dated tree sequences using three approaches. First, we inferred and dated a tree sequence using only the modern samples and evaluated the accuracy of the resulting allele age estimates. Second, we used these estimated allele ages to reinfer tree sequences (still only using modern samples) and evaluated the result. Third, we again used the allele age estimates to reinfer tree sequences, but constrained the estimates with progressively larger numbers of ancient samples. Allele age estimation accuracy was evaluated by comparing the true allele ages from the simulated tree sequence to the estimated allele ages using root mean squared log error (RMSLE) and Spearman's $\rho$. Allele ages were determined as described in the previous section. While the initial iteration step without ancient samples increases allele age estimation accuracy, adding samples drawn from empirical distribution of ancient sample ages does not have a noticeable effect on overall accuracy. However, increasing numbers of samples from sampling schemes (2) and (3) consistently improve both MSLE and Spearman's rank correlation. This is likely because older samples will "correct" larger numbers of derived alleles that are initially estimated to be too young.

#### S2.6 Archaic Descent Simulation

Tree sequences encode the genetic relationships between modern and ancient samples at every point in the genome. We used a published simulation model approximating the Out of Africa Event with archaic introgression into the ancestors of Eurasians and Papuans[16] to evaluate how well our inferred tree sequences reflect these relationships. Using an implementation of the simulation model from `stdpopsim`[8], we simulated 10 replicates of 15 Mb from chromosome 20, sampling 200 chromosomes each from the African, West Eurasian, East Asian, and Papuan populations. We slightly modified the ancient sampling scheme from the model by sampling three archaic individuals: the Denisovan (two chromosomes from the sampled Denisovan lineage, 2,203 generations ago), the Vindija (two chromosomes from the introgressing Neanderthal lineage, 1,725 generations ago), and the Altai Neanderthal (two chromosomes from the ancestral Neanderthal lineage, 3,793 generations ago). See Jacobs et al. (2019) Fig. S5 for a schematic summarising the model. This model defines a generation to be 29 years (compared to 25 years in our other analyses).

We used migration records from the simulated tree sequences to determine patterns of archaic local ancestry. This provided a "ground truth" set of introgressed spans of sequence in modern samples. Migrations from Denisovan and Neanderthal lineages to the ancestors of modern, non-African samples in the simulated trees provide the locations of these introgressed spans of sequence in each modern sample. Importantly, in this simulation setting introgressed tracts exist regardless of whether or not they carry derived alleles and thus may be undetectable.

We examined how well patterns of common ancestry between *sampled* archaic and modern individuals reflect archaic introgression. The simulation from Jacobs et al. (2019) modelled introgression as "pulses" of admixture from archaics to moderns at discrete points in time. Consequently, the time to most recent common ancestor (TMRCA) between a modern and archaic sample is only observed to be more recent than $T_{ArchaicModern}$, the split time of modern and archaic lineages occurring 20,225 generations ago, at marginal trees where introgression occurred in the ancestors of that modern sample. However, the TMRCA between sampled archaics and moderns may not be more recent than $T_{ArchaicModern}$ at all introgressed tracts: the TMRCA is older than the split time at marginal trees where the sampled and introgressing archaics are distantly related. TMRCAs that are more recent than $T_{ArchaicModern}$, but older than $T_{DenNea}$, the split time of Neanderthals and Denisovans occurring 15,090 generations ago, can be attributed to introgression from *either* Neanderthals or Denisovans. Signals of introgression from these two populations may be disentangled at marginal trees where the TMRCA of sampled archaics and moderns is more recent than $T_{DenNea}$.

We record tracts of common ancestry in this final category as well as the full set of introgressed tracts from the migration records using $n$x$c$ matrices, where $n$ is the sample size of modern individuals and $c$ is the number of 1 kilobase (kb) chunks in the simulated chromosome. Two introgression matrices were constructed, one each for the introgressing Denisovan and Neanderthal populations. At each cell in an introgression matrix, a "1" indicates introgression from the relevant archaic population in a modern individual in any marginal tree overlapping the 1 kb chunk. Three common ancestry matrices were constructed, one each for the Vindija, Altai, and Denisovan samples. In each common ancestry matrix, a "1" indicates that the TMRCA of the modern individual and *either* of the archaic individual's sampled chromosomes at any tree overlapping the chunk is less than $T_{DenNea}$.

Since TMRCAs of moderns and archaics more recent than $T_{DenNea}$ reflect introgression from Denisovan or Neanderthal lineages, values of "1" in the common ancestry matrices are guaranteed to only exist where "1"'s are observed in the corresponding introgression matrix. Common ancestry between moderns and the Denisovan more recent than $T_{DenNea}$ encompasses $\geq$ 99.9% of introgressed Denisovan tracts (Std Dev 0.04%). This is likely attributable to the simulated population size of only $N_e = 100$ in the introgressing Denisovan D1 and D2 lineages from Jacobs et al. (2019). The Vindija Neanderthal shares common ancestry with humans more recently than $T_{DenNea}$ for 61.3% (Std Dev 4.07%) of introgressed Neanderthal tracts. For the Altai, the value is 55.0% (Std Dev 1.99%).

Next, to investigate how well our inferred tree sequences reflect these observed patterns of introgression and common ancestry, we ran our iterative approach to infer a tree sequence of modern and archaic samples from the simulated data. We used an $N_e$ value of 10,000 and a mutation rate of $10^{-8}$ per base pair per generation when running `tsdate`. We then examined the inferred tree sequences, noting the spans of genome where the two "proxy ancestors" associated with each archaic individual have direct descendants among modern samples.

Observed descent from proxy archaic ancestors should overlap with signals of introgression where sampled archaic haplotypes are a better approximation of ancestral material than `tsinfer`'s inferred ancestors at a given epoch. This approach enables an understanding of the relationships between sampled archaic and modern individuals without needing to identify a split time between moderns and archaics. Results are expected to be heavily influenced by such factors as the sampling times of ancient individuals, relationships between modern and ancient populations, error rates, and the quality of ancestors estimated by `tsinfer`. Relationships between sampled and introgressing ancient lineages may particularly affect the amount of introgressed material that can be recovered from the inferred tree sequences since, as noted previously, in this simulation model more recent common ancestry is observed between the sampled and introgressing Denisovans than between the sampled and introgressing Neanderthals.

We used the inferred tree sequences to construct three additional $n$x$c$ matrices that record direct descent from proxy ancestors associated with the three sampled archaic individuals. In each cell in a matrix, a "1" records where a modern individual descends from the relevant proxy archaic sample in a marginal tree overlapping the 1 kb chunk. Comparing the binary matrices recording ground truth tracts of introgression or common ancestry from the simulations with the matrices recording inferred descent from archaic samples yielded true positives (TP), corresponding to introgression or common ancestry in the simulation and descent in inferred tree sequence, false positives (FP), where no introgression or common ancestry is noted in the simulation but descent is inferred, and false negatives (FN), where introgression or common ancestry is noted in the simulation but no descent is inferred. Table S1 shows the resulting rates of precision and recall, which are defined as follows:

$$Precision = \frac{TP}{TP + FP}, \tag{4}$$

$$Recall = \frac{TP}{TP + FN}. \tag{5}$$

The proportion of all modern genetic material that is introgressed from each archaic population, the proportion of modern genetic material sharing common ancestry with archaic samples more recently than $T_{DenNea}$, and the proportion of modern genetic material that is inferred to descend from archaic samples are also given in Table S1.

The simulation results indicate that descent from the Vindija and Denisovan, and to a lesser extent the Altai, recovers ground truth introgression and shared ancestry with high precision. In the case of the Neanderthals, where shared ancestry tracts incompletely represent introgressed tracts, inferred descent recovers more shared ancestry tracts than introgressed tracts.

It is possible to increase recall by examining patterns of common ancestry between sampled archaics and moderns occurring more recently than a given split time in the inferred tree sequence, as was performed in the simulated tree sequence. The precision and recall for recovering ground truth introgressed and shared ancestry tracts at different split times is analysed in detail in Fig. S5, where observed precision and recall values are obtained by testing split times in intervals between the time of each archaic sample and $T_{DenNea}$. These results show that progressively older split times recover nearly all shared ancestry tracts, though at lower levels of precision. While it is possible to use these results to choose time cutoffs for each sample which maximise recall at a given level of precision, the optimal values derived from simulations will doubtlessly differ from the real data due to the imperfect modelling of demographic history as well as varying levels of error. In this study we chose to report tracts that directly descend from these samples in order to avoid choosing a cutoff time while likely maximising precision.

| | Vindija | | Altai | | Denisovan | |
|---|---|---|---|---|---|---|
| | Introgressed | Common Ancestry | Introgressed | Common Ancestry | Introgressed | Common Ancestry |
| Precision | 0.92 (0.03) | 0.87 (0.05) | 0.69 (0.08) | 0.59 (0.08) | 0.86 (0.04) | 0.86 (0.04) |
| Recall | 0.18 (0.05) | 0.28 (0.07) | 0.42 (0.04) | 0.65 (0.06) | 0.61 (0.11) | 0.61 (0.11) |
| Simulated % of moderns | 2.86% (1.12%) | 1.77% (0.71%) | 2.86% (1.12%) | 1.57% (0.60%) | 0.99% (0.24%) | 0.98% (0.24%) |
| Inferred % of moderns | 0.53% (0.15%) | | 1.65% (0.31%) | | 0.69% (0.12%) | |

Table S1: Results of archaic descent simulations evaluating how well direct descent from proxy archaic haplotypes in inferred tree sequences recovers ground truth introgressed tracts of sequence as well as tracts where modern samples and sampled archaics share common ancestry more recently than $T_{DenNea}$. Definitions of precision and recall are given in Equations 4 and 5. "Simulated % of moderns" refers to the percentage of genetic material from all modern individuals in each simulation that is introgressed from archaics or shares recent common ancestry with archaics. "Descendant % in moderns" is the percentage of modern genetic material in each inferred tree sequence which descends from each archaic individual. In each cell, the mean values from 10 simulation replicates is given with the standard deviation in brackets.

### S3 Dataset Preparation

#### S3.1 Downloading and preparing publicly available data

The TGP Phase 3 GRCh37 VCF was accessed from TGP release 20130502[17], while the TGP GRCh38 variant calls were accessed from the European Variation Archive under accession number ERZ822766[18]. SGDP data was downloaded from `https://sharehost.hms.harvard.edu/genetics/reich_lab/sgdp/phased_data/PS2_multisample_public`[19]. The HGDP statistically phased dataset was downloaded from `ftp://ngs.sanger.ac.uk/production/hgdp/hgdp_wgs.20190516/statphase/`[20]. Note that HGDP does not provide phased data from the X chromosome. The Reich laboratory provides a compilation of 3589 genotyped ancient individuals available at `https://reich.hms.harvard.edu/downloadable-genotypes-present-day-and-ancient-dna-data-compiled-published-papers` (the full list of citations for these papers is available from this link). The sequenced Denisovan, Vindija, Altai, Ust-Ishim, Loshbour, and LBK-Stuttgart samples were downloaded from `http://cdna.eva.mpg.de/neandertal/Vindija/VCF/` and the Chagyrskaya Neanderthal from `http://ftp.eva.mpg.de/neandertal/Chagyrskaya/`.

The GRCh38 reference assembly was used in all analyses with the exception of the TGP variant age estimation analysis where GRCh37 was used. Only a GRCh37 version of the SGDP and ancient datasets were available, so these datasets were lifted over to GRCh38 using the Picard tool[21] and the hg19 to hg38 UCSC LiftOver chain[22].

The four archaic individuals are phased using Beagle 5.1 without a reference panel and without imputation[23]. The lack of a suitable reference panel likely results in extensive phasing error, but the high levels of homozygosity in archaic genomes mean that errors will only occur between heterozygous sites. Furthermore, errors will only effect downstream analyses when a descendant haplotype crosses a switch error. Indeed, these errors seem to have a limited effect, as running the inference pipeline with unphased archaic genomes does not change any of the conclusions we draw.

#### S3.2 Afanasievo Family

We generated between 5 to 8 lanes of shotgun sequencing data using an Illumina HiSeqX10 instrument (2 x 101 cycles and reading out the indices with 2 x 7 cycles) from four individuals from the Afanasievo culture, using sequencing libraries for which in-solution enrichment data was previously published in Narasimhan et al. Science 2019 (Supplementary Table 2). We merged the paired sequences and mapped to the human genome reference sequence using the samse command in BWA v.0.7.15-r114053 with the parameters -n 0.01, -o 2, and -l 16500, and removed duplicated molecules as described in the original publication that generated in-solution enrichment data on these libraries. The average coverage measured on the autosomal SNP targets used for the in-solution enrichment experiment was 10.8x for I3388 (the mother in the family), 25.8x for I3950 (the father in the family), 21.2x for I6714 (one of the sons in the family), and 25.3x for I3949 (the other son of the family).

The Afanasievo "quartet" allows for reliable phasing of ancient genomes. We used bcftools (v1.10.2) to calculate genotype likelihoods of biallelic SNPs in the Afanasievo samples. We used Beagle 4.0[23] by providing a genotype likelihood file (gl argument) and pedigree file (ped argument) to phase this family without any reference panel and imputation. We excluded sites with $\max(GP) < 0.99$.

#### S3.3 Inferring a unified genealogy

Once VCF files from each dataset were prepared, we created `.samples` files, a file format used with `tsinfer`, for each constituent dataset using all biallelic SNPs with high-confidence ancestral alleles from Ensembl release 100[24]. The `.samples` files were then combined using the `merge` functionality in `tsinfer`, where the genotypes of variants missing from at least one sample were marked as "missing data" to be imputed by `tsinfer`. A tree sequence was then inferred and dated for each autosome using modern samples from the HGDP, SGDP, and TGP datasets. Empirical lower bounds on allele age were gathered from all ancient samples (including unphased and genotyped samples). Variants present in an ancient individual but missing in all modern datasets were not used. Ancient lower bounds at each variant site were compared with estimates from the inferred tree sequence of modern samples. The estimated age of a site in the inferred tree sequence is defined as the oldest most recent common ancestor which possesses a derived variant at each site. At sites where the empirical lower bound from ancient samples is older than our estimated age, this bound was used as the variant age. With these constrained age estimates, the tree sequence was re-inferred using the four archaic individuals and the Afanasievo quartet. Details of this process can be found at `https://github.com/awohns/unified_genealogy_paper/all-data`.

154 modern individuals appear in more than one dataset. This consists of 130 individuals which appear in both SGDP and HGDP, as well as 24 individuals which appear in both SGDP and the TGP GRCh38 release. Duplicated individuals are retained in all analyses for the following reasons: the scalability of our methods mean that removal of these samples would result in negligible savings in computational resources, none of the analyses we conducted would be adversely affected by duplicated samples, and inspecting the duplicated samples revealed disagreements in genotyping and phasing. Across all 154 duplicated samples on chromosome 20, the mean proportion of genotype calls that are incompatible is 1.87% (standard deviation 1.32%). This figure was found by examining the variant sites shared between each pair of duplicated individuals and computing what proportion of these variants had differing numbers of derived alleles. This discrepancy can be explained by the fact that some of the duplicated samples were sequenced using different libraries, that different variant calling pipelines were used, by the imputation of missing genotypes in HGDP individuals, and also potentially by the effects of lifting over SGDP individuals from GRCh37 to GRCh38. We determined phasing mismatch by examining the heterozygous sites in duplicated individuals. This resulted in a "mismatch score" for each pair, calculated as the percentage of derived allele mismatches between the pairs at heterzygous sites. Two mismatch scores are calculated for each pair, based on the two possible phasing alignments. At each duplicated pair, we find the lower mismatch score. The mean across all pairs is 46.8% (standard deviation 2.5%), indicating that there is almost no consistency in chromosome-wide phasing between the duplicated pairs. Nevertheless, in the unified tree sequence we find that individuals share a parent node with one another over 61.5% of the tree sequence (standard deviation 9.9%).

### S4 Pairwise Time to Most Recent Common Ancestor

We use the unified and dated tree sequence to calculate the pairwise TMRCA for sampled haplotypes from all 215 populations in the combined tree sequence of HGDP, SGDP, TGP and ancient samples. This was accomplished by iterating over the trees in the tree sequence and at each tree calculating the TMRCA between pairs of chromosomes using the `mrca` function implemented in `tskit`. For efficiency, we down-sampled the chromosomes by randomly selecting up to ten samples from each population. In populations with fewer than 10 samples, all samples are used. We then find the weighted logarithmic average of all TMRCAs associated with each population combination, weighted by the span of the tree associated with each TMRCA. The logarithmic average is used as we expect the variance of TMRCAs to increase substantially with age: all date values are log transformed before analysis for this reason. The constrained age estimates produced by `tsdate` were used in this analysis (see Methods).

The histograms of TMRCAs for each population combination can be found in Supplementary Interactive Fig. 1. Three of these histograms are also shown in Fig. 2.

### S5 Empirical Estimates of Thousand Genome Project Variant Ages on Chromosome 20

We compared the estimated ages of GRCh37 chromosome 20 TGP Variants from `tsdate`, `Relate` and `GEVA`. We used `tsinfer` to infer a tree sequence of TGP individuals with the GRCh37 recombination map, mismatch ratios of 1, and a precision setting of 15. The tree sequence was dated using `tsdate` with $N_e$ set to 10,000 and a mutation rate of $10^{-8}$ mutations per base pair per generation. Default settings were used for all other parameters. Allele ages were estimated by using the arithmetic mean of the nodes above and below the oldest mutation. Allele age estimates from `GEVA` were gathered from `https://human.genome.dating/download/index`. We used the mean allele age estimate from the joint clock as a point estimate of allele age and the upper limit of the 95% confidence interval from the joint clock as an upper bound estimate[12]. `Relate` TGP allele age estimates were downloaded from `https://zenodo.org/record/3234689#.X8EXyapKhTZ`. Since the publicly available `Relate` age estimates are calculated separately in each TGP population, we averaged age estimates and upper bounds for alleles appearing in more than one population. The point estimate of allele age was obtained by taking the mean of the upper and lower age bounds.

Empirical lower bounds were provided by radiocarbon dates of ancient samples. These dates were derived from the Reich Laboratory dataset for all ancient individuals with the exception of the Afanasievo family (which we assigned to 4.6 kya), Chagyrskyaya Neanderthal (Chagyrskaya 8) (80 kya)[25], Vindija Neanderthal (Vindija 33.19) (50 kya)[26], Altai Neanderthal (Denisova 5) (110 kya), Denisovan (Denisova 3) (63.9 kya)[27], Ust'-Ishim (45 kya)[28], Loshbour (8 kya)[29] and LBK (7 kya)[29]. If multiple ancient samples carried a derived allele, the oldest sample carrying the allele was used as the lower bound on the age of that allele. This resulted in a combined dataset of 3,734 ancient individuals.

A set of 659,804 sites for which age estimates can be found from all three methods was assembled. This excluded singletons and $n-1$ tons, which `GEVA` did not estimate, sites where the derived and alternate alleles were inconsistent (as `GEVA` estimates the age of alternate alleles), indels, and sites with low quality ancestral states. The relationship of allele age estimates from each method to allele frequency is shown in Fig. S6, the distribution of ages is shown in Fig. S7, and the comparisons of allele ages from the differing methods to one another is shown in Fig. S8. The combined set of ancient samples carried derived alleles at 76,889 of these sites. At each site, the oldest ancient sample carrying a derived allele was used as the empirical lower bound on the age of the site. A comparison of the estimated allele ages from the three methods with these bounds is shown in Fig. 3a.

It is important to note that the distribution of age estimates varies between the three methods (see Fig. S7), with `Relate` showing a higher mean estimated age compared to the other two methods. The `tsdate` mean estimated allele age at the 659,804 comparable sites is 5,919 generations and the mean upper bound on allele age is 9,012 generations. For `Relate`, the equivalent values are 6,816 and 11,732 generations. Since `Relate` provides an estimate for each population, these values were first found by averaging the available population-based estimates for each site, then averaging all the sites. For `GEVA` the equivalent values are 5,192 and 6,147 generations.

### S6 Descent from Ancient Individuals

We used the unified, inferred tree sequence to investigate the genealogical relationships of ancient and modern samples. A number of studies have sought to infer introgressed haplotypes from archaic individuals[30–33] and one has inferred an ancestral recombination graph from modern and archaic individuals[34]. As shown in S2.6, our approach identifies tracts of introgressed sequence, particularly regions where sampled and introgressing archaic individuals share more recent common ancestry, and requires no assumptions about the nature of introgression.

We evaluate descent from the Afanasievo family, Denisovan, and Vindija Neanderthal among younger samples in Fig. 3 and Extended Data Figs. 6-8. The simplified tree sequence of 3,754 individuals on chromosome 20 (with unary nodes retained) is used for this analysis. Regions of over 1 Mb without variant sites were trimmed from the inferred tree sequence, as well as regions before the first variant site and after the last variant site. We establish descent using the *proxy nodes* associated with each ancient sample in the inferred tree sequence, as detailed in the Methods section.

Descent from ancient individuals is described using two statistics. First, the genomic descent statistic defined in Scheib et al. 2019 is used to evaluate the overall amount of genetic material in each population which descends from the proxy ancestors associated with ancient individuals. The results of this analysis are shown for the Denisovan (Fig. 3b and Extended Data Fig. 7), Afanasievo (Extended Data Fig. 6a), and Vindija (Extended Data Fig. 8). Second, we split chromosomes into 1 kb chunks and assess descent from the ancient individuals in each chunk, as described in section S2.6. Pearson product-moment correlation coefficients were calculated using `Numpy`[35] for the $n$ x $c$ matrices of descent, where $n$ is the number of descendants and $c$ is the number of 1 kb blocks. The correlation coefficients were then hierarchically clustered using the UPGMA algorithm implemented in `Scipy`[36]. The results of this analysis are shown for the Denisovan (Fig. 3c) and Afanasievo (Extended Data Fig. 6b).

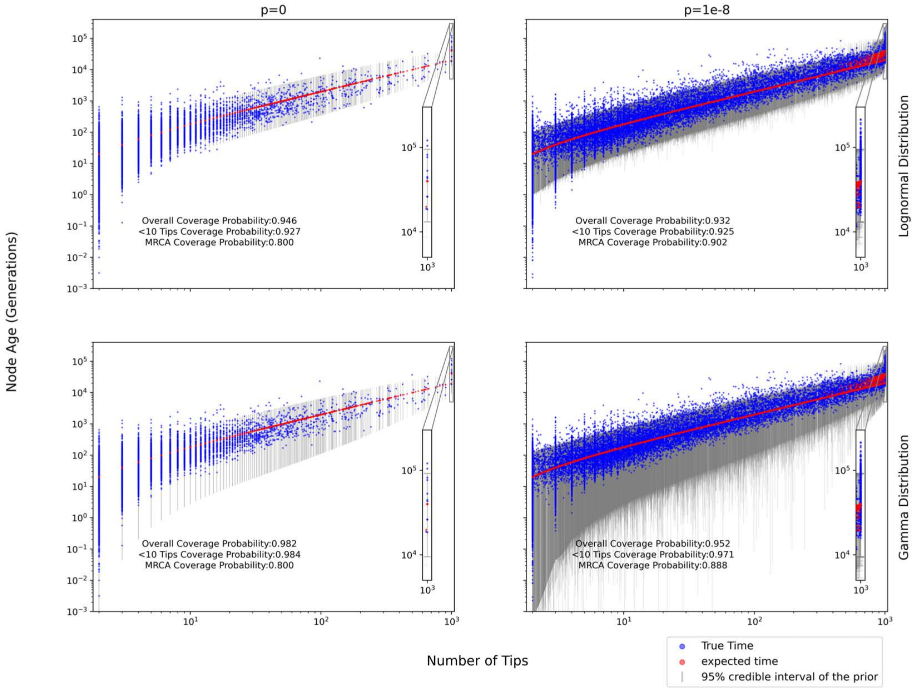


Figure S1: Accuracy of the `tsdate` node age prior distribution. The subplots in the left column show node ages from ten `msprime` simulations of length 500 kb with 1000 samples, $N_e$ =10,000, and $r = 0$. The right column shows shows results of ten simulations using $r = 10^{-8}$ and the same parameters otherwise. The top plots show the accuracy of the lognormal approximation to the conditional coalescent and the bottom show the gamma approximation.

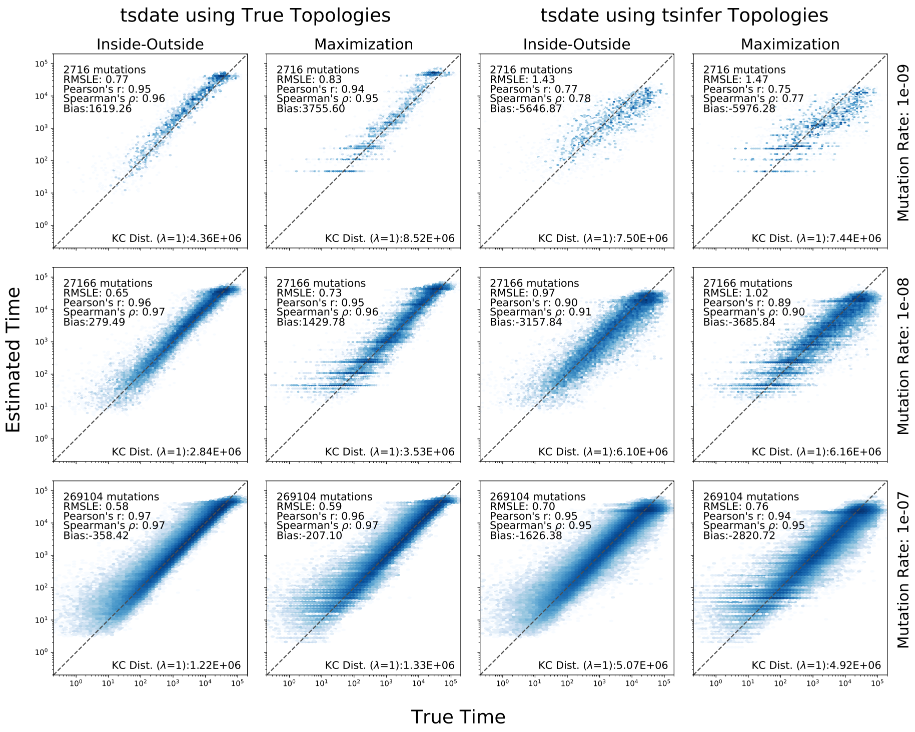


Figure S2: Accuracy of `tsdate` at various mutation rate settings. Each row shows the results of ten `msprime` simulations of 1 Mb with 500 samples, $N_e$= 10,000, and $r = 10^{-8}$. Three different values of $\mu$ were used, $10^{-9}$, $10^{-8}$, and $10^{-7}$. In each subplot simulated allele ages are on the x-axis and estimated allele ages are on the y-axis. The first two columns show the results of running `tsdate` on topologies simulated by `msprime`. The third and fourth columns show the results when running `tsdate` on topologies inferred by `tsinfer` from the simulated genotype data. The results of the inside pass followed by either an outside pass or outside-maximisation pass are shown.

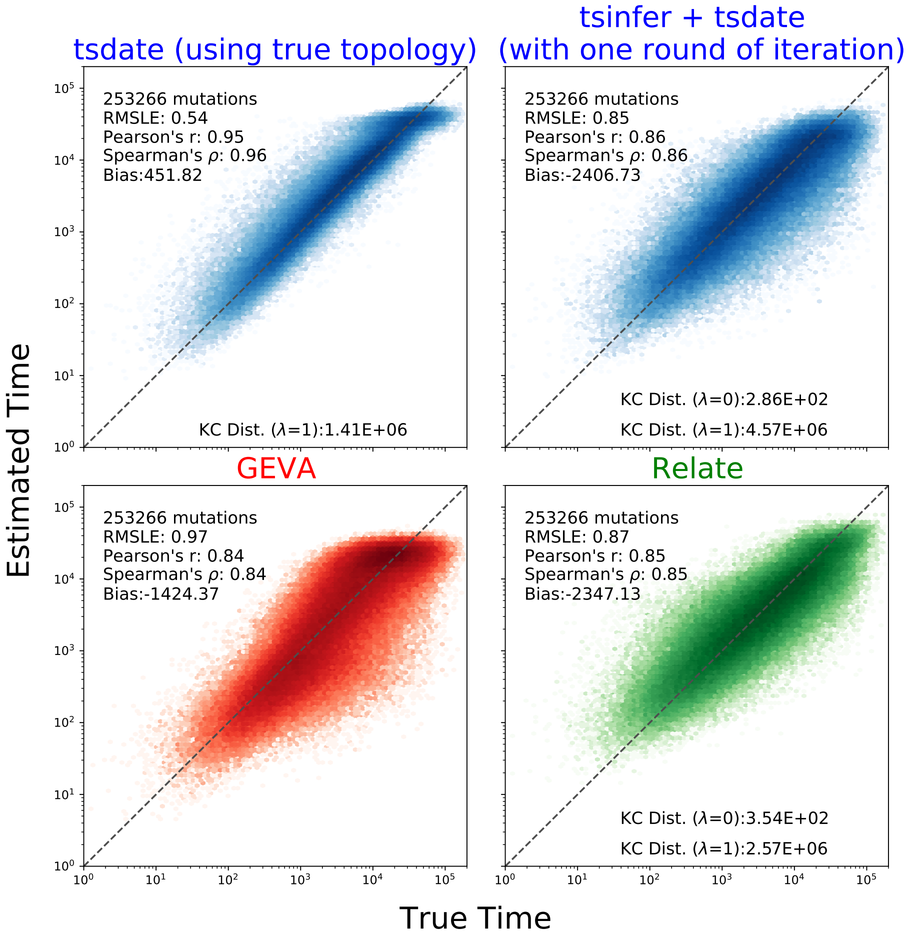


Figure S3: Comparison of the accuracy of `tsdate`, `GEVA`, and `Relate` on simulated tree sequence topologies. Thirty `msprime` simulations with 250 samples, 5 Mb of sequence, $N_e = 10{,}000$, and $\mu = r = 10^{-8}$ were used. The x-axis in all plots is the age of derived alleles from the simulation; estimated allele ages from the labeled method are on the y-axis. Top left subplot shows derived allele age estimates from running `tsdate` on simulated topologies. Top right subplot shows results from inferring a tree sequence topology with `tsinfer`, dating the tree sequence with `tsdate`, and using the resulting date estimates to reinfer and redate the tree sequence. Lower subplots show allele age estimates from `GEVA` and `Relate`. Only alleles dated by all methods are shown: this excludes singletons, $n-1$ tons, and alleles that were deemed to map poorly by `tsdate` or `Relate`.

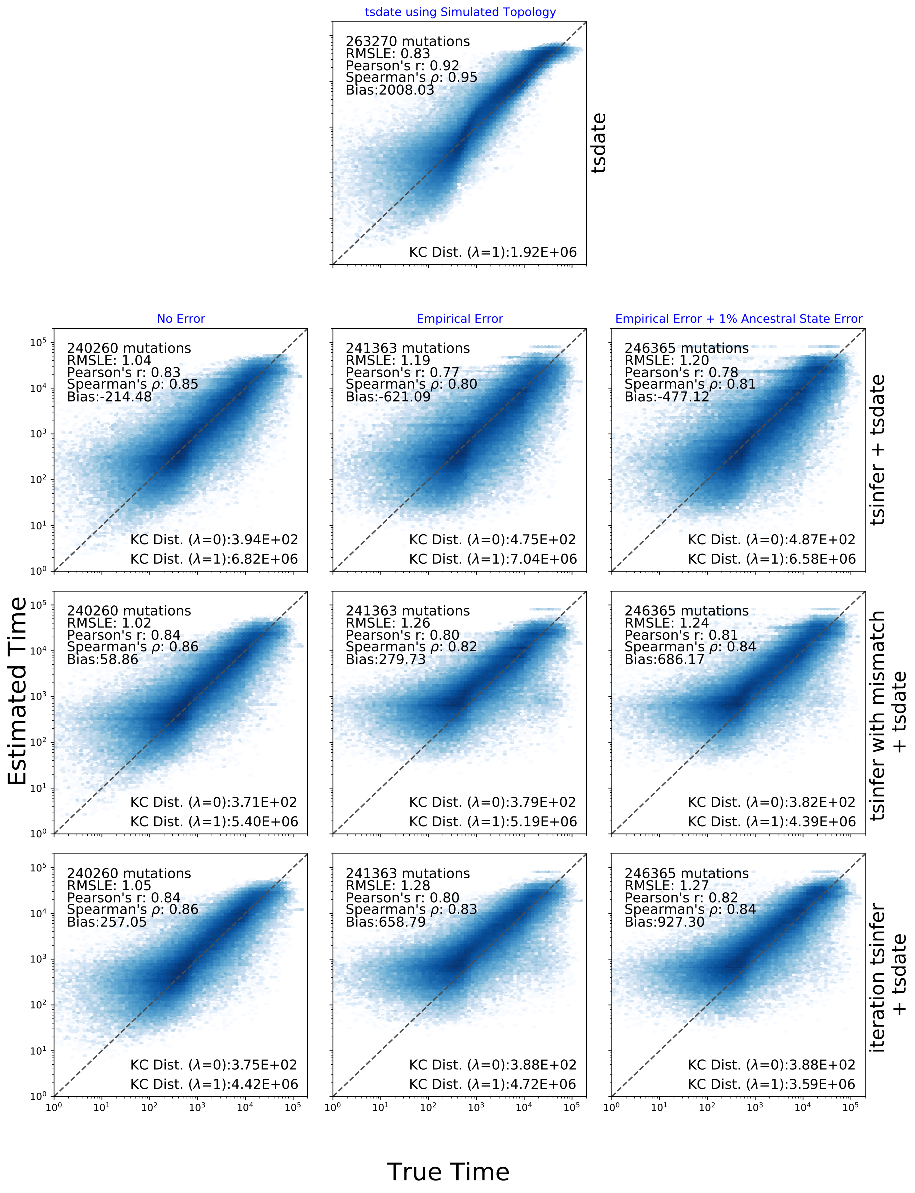


Figure S4: Evaluation of `tsdate` on simulated chromosome 20 data. `stdpopsim` and `msprime` were used to simulate human chromosome 20 using the three population Out of Africa model[7]. 100 samples from YRI, CEU, and CHB were used. The top row shows the results of using `tsdate` on the simulated tree sequence topology. The second row shows the accuracy of `tsdate` on topologies from `tsinfer`. The third row shows the results of using topologies produced using `tsinfer`'s full Li and Stephens model with error. The final row shows the results of feeding the inferred dates from row two back into `tsinfer` and running `tsdate`. The first column in rows 2-4 shows simulations without genotype error, the second shows simulations with an empirical genotype error model and the third shows an empirical genotype error model and 1% ancestral state assignment error.

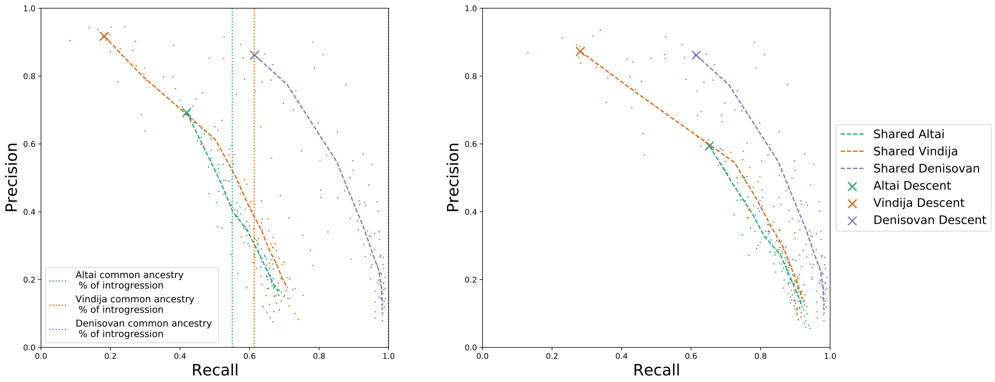


(a) Introgressed Tracts

(b) Shared Ancestry Tracts

Figure S5: Precision-recall curves for recovering relationships between archaic and modern samples using inferred tree sequences. (a) Accuracy of recovering introgressed tracts from archaic populations. Dotted vertical lines indicate total amount of modern genetic material where modern samples and each archaic individual share common ancestry in the *simulated* tree sequence more recently than $T_{DenNea}$ as a proportion of total introgressed material. Common ancestry in the *inferred* tree sequences occurs when the TMRCA of modern samples and an archaic sample occurs more recently than a specified time cutoff. Cutoffs ranging from the time of each archaic sample to $T_{DenNea}$ are tested, scatter plots show the results from each archaic sample at each cutoff and each simulation replicate. Dashed lines show the average precision and recall across replicates at each cutoff. Older time cutoffs result in progressively lower precision and higher recall. Precision is maximised at points marked with an “X”, where the cutoff is equal to the time of the sample, i.e. where only material directly descending from archaic proxy samples is used. (b) Precision-recall curves for recovering only the shared ancestry tracts for the three sampled archaics indicated by the dotted vertical lines in (a).

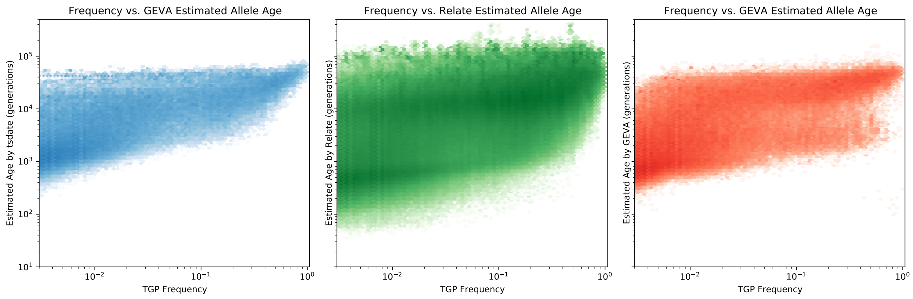


Figure S6: Estimated age of TGP variants from `tsdate`, `GEVA` and `Relate` compared to allele frequency. Estimates of allele age are found using the arithmetic mean of the node ages above and below the mutation for `tsdate` and `Relate`. For `GEVA`, the mean age of the joint clock estimate is used. The same set of alleles is also used in Figs. 3a, S7, and S8

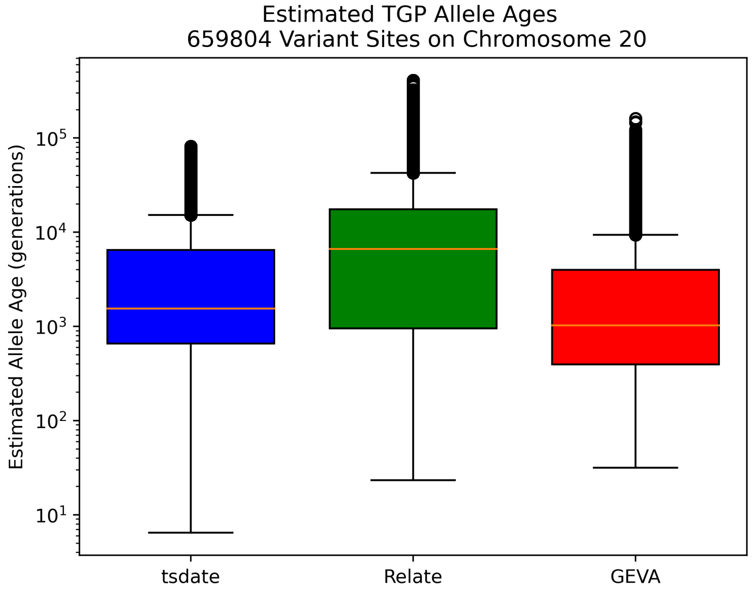


Figure S7: Average age of TGP variants from `tsdate`, `GEVA` and `Relate`. Box plot elements are defined in Fig. 6a. Note, the age range of `tsdate` depends on the number of time slices specified by the user (default settings were used for this paper).

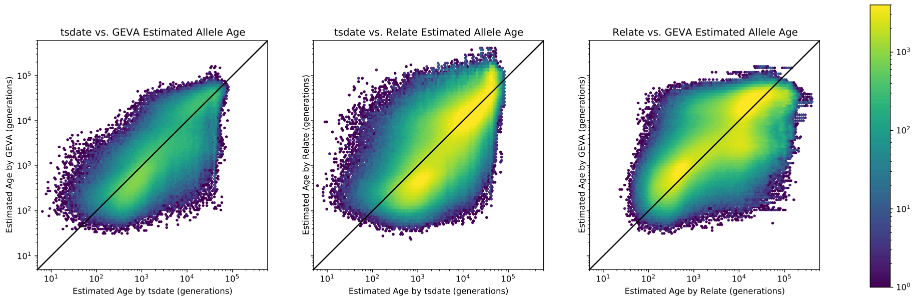


Figure S8: Comparison of TGP derived allele age estimates from `tsdate`, `GEVA` and `Relate`. Each hexbin plot shows estimates of variant age by two different methods.

#### Supplementary Video and Interactive Figure

Supplementary Video 1 is available at: `https://www.youtube.com/watch?v=AvV0zBSdxsQ`. The video shows the estimated geographic locations of ancestors of HGDP, SGDP, Neanderthal, Denisovan, and Afanasievo samples over time. Each dot represents an edge in the tree sequence of chromosome 20, where the time and geographic location of the parent and child nodes of the edge have been estimated. The locations of edges at each point in time are plotted using the great circle between the parent and child nodes. Edges are coloured by the region of the descendants of the child node. If an ancestral lineage has ancestors in multiple regions, its colour is the average of the respective colours of each region.

Supplementary Interactive Figure 1 is available at: `https://awohns.github.io/unified_genealogy/interactive_figure.html`. This dynamic version of Fig. 2 shows the span-weighted TMRCA histogram relating each pair of the 215 populations in the integrated tree sequence of chromosome 20. Hovering over each cell displays the relevant histogram, with a vertical bar indicating the logarithmic mean TMRCA.
